# Global analysis of cancer cell responses to USP9X inhibition

**DOI:** 10.1101/2025.06.18.660475

**Authors:** Philipp Schenk, Shane M. Devine, Simon A. Cobbold, Ching-Seng Ang, Niall D. Geoghegan, Dale J. Calleja, Dylan H. Multari, Vineet Vaibhav, Bernadine G.C. Lu, Theresa A. Klemm, Laura F. Dagley, Kym N. Lowes, Nicholas Williamson, Pieter J. A. Eichhorn, Ashley P. Ng, Rebecca Feltham, David Komander

## Abstract

The ubiquitin specific protease (USP) enzyme USP9X is amongst the best studied human deubiquitinases (DUBs), with a myriad of described targets and cellular roles. In cancer, USP9X has been touted as both an oncogene and a tumour suppressor in different contexts, which has confounded the field and questioned its viability as a cancer target. We here describe WEHI-092, a novel piperazine-based USP9X specific small molecule inhibitor and map its binding site to a unique region in the USP9X fingers subdomain, distinct from known DUB inhibitor binding sites. Using proteomics and ubiquitinomics, we show that USP9X has a distinct set of substrates compared to USP7 indicating remarkable DUB target specificity, yet the substrate profile of USP9X varies significantly across cancer cell lines. Interestingly, we reveal a core set of 17 proteins commonly regulated by USP9X in most or all cell lines, which we consider as proximal biomarkers for USP9X inhibition. Consistent with our proteomic analyses, we show that WEHI-092 treatment arrests cells in metaphase without inducing cell death, which may account for growth suppression seen in long-term clonogenic assays in most cancer cell lines, and positions USP9X inhibitors as a new potential class of selective mitotic poisons.

## Introduction

Protein ubiquitination is a key regulator of protein homeostasis, and as such has many recognised roles in human disease settings, in particular cancer (Deng *et al*, 2020). The attachment of ubiquitin to proteins most commonly triggers their degradation through the ubiquitin proteasome system (UPS), which facilitates the clearance of old and damaged proteins but can also regulate cellular cascades such as the cell cycle (Hershko, 1997). Overall, more than 5% of the human genome encodes for proteins that attach, bind, or remove a vast array of distinct and dynamic ubiquitin modifications (Agrata & Komander, 2025). Within these proteins, more than 700 E3 ligases facilitate ubiquitination in conjunction with E1 activating and E2 conjugating enzymes, while ∼100 deubiquitinases (DUBs) remove ubiquitin signals from proteins (Clague *et al*, 2019).

Many oncogenes are under the control of the UPS, and are ubiquitinated and degraded, often in a highly regulated fashion. The UPS is exploited pharmacologically, most prominently via targeted protein degradation (TPD) approaches using proteolysis-targeting chimeras (PROTACs) or molecular glues (Békés *et al*, 2022; Zhao *et al*, 2022). Some of the most desired cancer targets, however, remain undruggable due to a lack of efficacious small molecule binders. An orthogonal way to drive an oncoprotein towards degradation, can be achieved through inhibition of the DUB that stabilises the oncoprotein (Harrigan *et al*, 2018). Indeed, hundreds of papers report deregulated DUB activity in cancer, where DUB deregulation stabilises oncoproteins, or destabilises tumour suppressors (Dewson *et al*, 2023). Hence, DUB inhibitors provide an orthogonal means to induce oncoprotein degradation.

Ubiquitin specific protease (USP) DUBs have received most pharmacological attention. USP DUBs are typically promiscuous regarding the ubiquitin signals they remove but achieve target specificity via substrate-binding domains (Mevissen & Komander, 2016). The more than 50 human enzymes in the family comprise several known oncogenes (Dewson *et al*, 2023). However, drug discovery for USP DUBs has proven challenging, and truly specific USP inhibitors have only been reported in the last decade, for seven USP enzymes (reviewed in Liu *et al*, 2025; Schauer *et al*, 2020). A fascinating diversity of inhibitory mechanism has been revealed in structural studies (Kazi *et al*, 2025).

USP9X is amongst the best studied USP enzymes. More than 400 papers have focused on this enzyme, revealing roles in DNA damage response, cell cycle regulation, ribosomal quality control, endosomal trafficking, TGFβ and Hippo signalling, and others (reviewed in Murtaza *et al*, 2015; Gao *et al*, 2024). USP9X is essential for organism (especially brain) development (Pantaleon *et al*, 2001; Stegeman *et al*, 2013) as originally described in the *fat facets* phenotypes in drosophila (Fischer-Vize *et al*, 1992) and mouse (Wood *et al*, 1997). Recent papers have focused on the roles of USP9X in cancer, where conflicting literature has shown USP9X to act as an oncoprotein or as a tumour suppressor in different cancer types. An oncogenic role for USP9X has been reported broadly across many cancers, and since many putative USP9X substrates are interesting pharma targets (such as MCL-1, β-catenin, ERG), USP9X became a promising pharmaceutical target itself (Schwickart *et al*, 2010; Wang *et al*, 2014; Lu *et al*, 2019; Taya *et al*, 1999). Conversely, USP9X was reported as a frequently mutated gene in pancreatic ductal adenocarcinoma (PDAC) (Pérez-Mancera *et al*, 2012), and other human cancers (Cheasley *et al*, 2021; Hunter *et al*, 2015; Sisoudiya *et al*, 2023), suggesting tumour suppressor roles. Indeed, several putative USP9X substrates are tumour suppressors (LATS2, YAP1, FBW7) (Zhu *et al*, 2018; Li *et al*, 2018; Toloczko *et al*, 2017; Khan *et al*, 2018). While the tumour suppressor role of USP9X in PDAC has been challenged (Cox *et al*, 2014b; Pal *et al*, 2017; Liu *et al*, 2017), these contrasting roles questioned the validity of USP9X as a cancer target, and reinforced the need to define the molecular landscape and clinical context in which targeting USP9X would be most effective.

Some of the discrepancies in the literature arise from an undue focus on single-protein relationships (USP9X regulates target X), which was sometimes exaggerated by inappropriate tools. More than 30 papers have assigned the effects of Degrasyn/WP1130 and its derivatives G9/EOAI3402143 (Bartholomeusz *et al*, 2007a, 2007b) to USP9X inhibition, despite data that WP1130 is a non-specific DUB inhibitor (Ritorto *et al*, 2014; Kapuria *et al*, 2010; Peterson *et al*, 2015).

We recently reported FT709 as a first, highly specific USP9X inhibitor (Clancy *et al*, 2021). While its molecular mechanism of action has remained unclear, FT709 revealed a role for USP9X in ribosomal quality control in a cancer cell line (Clancy *et al*, 2021), but failed to confirm many previously reported USP9X substrates.

We here report WEHI-092 as a next generation USP9X inhibitor. The simplified chemical scaffold retains excellent USP9X specificity, which we structurally explain through Hydrogen Deuterium Exchange Mass Spectrometry (HDX-MS) and mutagenesis studies. WEHI-092 is highly potent in cells, and broadly arrests cell growth in a panel of cancer cell lines, and strikingly selective in inducing cell death in one of 54 cell lines. Unbiased ubiquitinomics and proteomics experiments across two and eight cell lines, respectively, unveil unprecedented insights into USP9X biology. We identify a set of high confidence USP9X-regulated proteins across all cell lines studied, representing potential biomarkers for USP9X inhibition. Surprisingly, USP9X inhibition regulates a distinct set of targets in each cell line, and such pleiotropic cell-type specific effects may explain discrepancies in the literature. Importantly, collective analysis of all proteomic data uncovers commonalities, showing that USP9X inhibition leads to mitotic metaphase arrest without cell death, explaining growth retardation phenotypes.

## Results

### Identification of a USP9X specific small molecule inhibitor

While the USP9X inhibitor FT709 demonstrated potency *in vitro* and *in cellulo*, it bears a challenging chemical structure with a central symmetric bicyclic core resulting in a complex synthesis route and high cost of production (Follows *et al*, 2019). During our search for DUB inhibitors within our chemical libraries, we identified WEHI-553 (**Fig. S1A**) as a compound that inhibited the purified catalytic domain of human USP9X (residues 1552-1970) *in vitro* with an IC50 of 4.2 µM in a ubiquitin-rhodamine (Ub-Rho) based screening campaign (Klemm *et al*, 2020) (**Fig. S1B**). WEHI-553 was structurally similar to FT709 but features a simpler and more chemically adaptable piperazine group as a central core. Structure-activity relationship (SAR) studies improved the IC50 towards USP9X > 16-fold with the best compound, WEHI-092 (**Fig. 1A, Fig. S1A**), displaying an IC50 of 254 nM, compared to 124 nM for FT709 (**Fig. 1B**). Direct compound binding to the USP9X catalytic domain was measured by surface plasmon resonance (SPR) and revealed a KD of 69 nM for FT709 vs. 1 µM for WEHI-092 (**Fig. 1C, Fig. S1C, D**). Incubation of the USP9X catalytic domain with 50 µM WEHI-092 inhibited cleavage of K48-linked diubiquitin (**Fig. 1D**) and prevented the modification of USP9X with ubiquitin propargylamide (Ub-PA), a suicide substrate (Ekkebus *et al*, 2013) that covalently modifies the catalytic Cys1566 of USP9X (**Fig. 1E**). Comparison with reported inhibitors for USP9X in this assay, showed no or rather weak effects of WP1130 and EOAI3402143 on the purified USP9X catalytic domain (**Fig. S1B, E**).

**Fig. 1.**
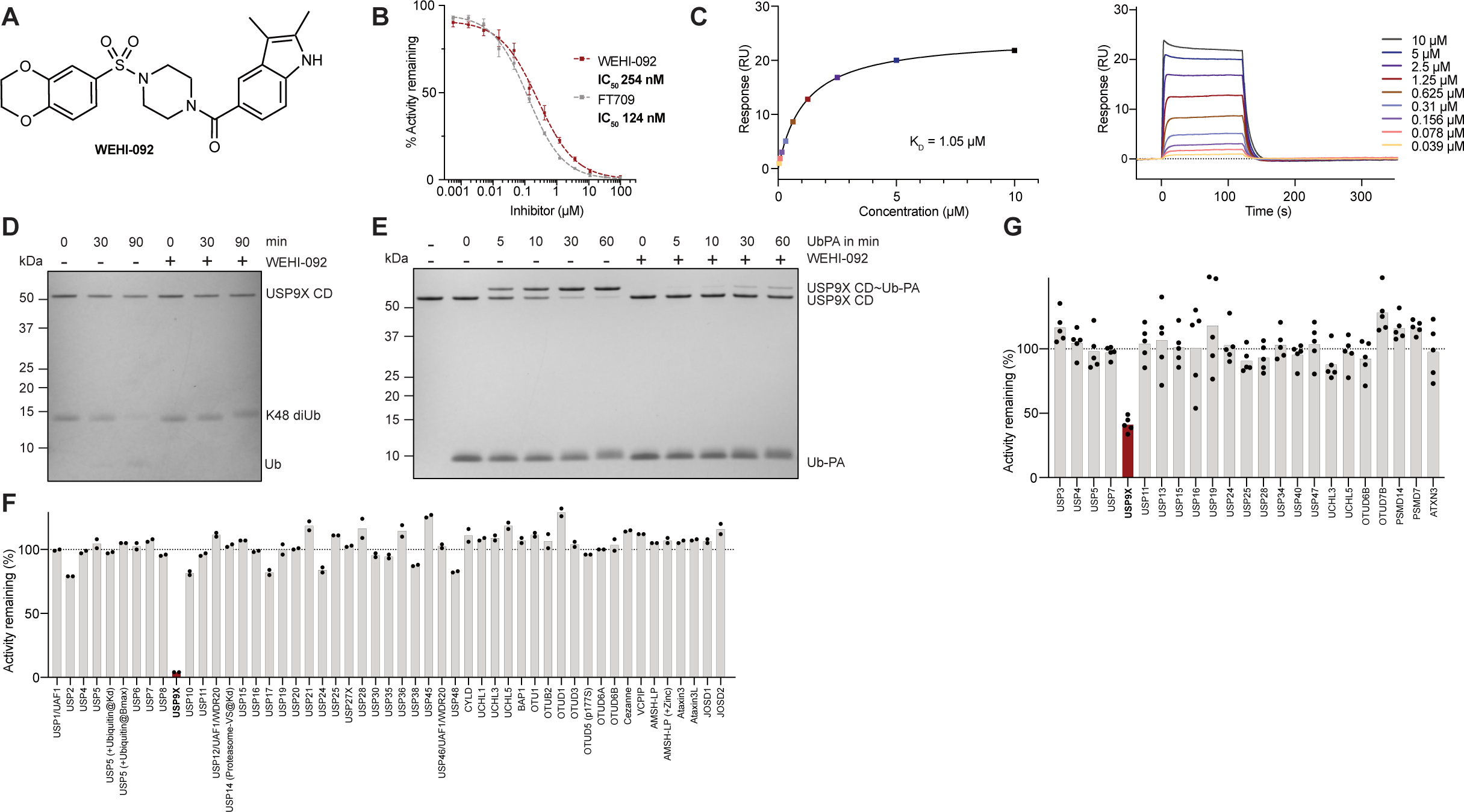
Identification of WEHI-092 as a novel inhibitor for the deubiquitinase USP9X. **(A)** Chemical structure of WEHI-092. (**B**) WEHI-092 and FT709 activity against recombinant USP9X catalytic domain determined from Ub-Rho cleavage assay. Data shown is the mean of four independent biological repeats with two technical replicates per experiment. Curve shown is the nonlinear curve fit generated using GraphPad Prism (v10.3) and was used for calculating the IC50 values. Error bars, SEM. (**C**) Direct binding of WEHI-092 to the USP9X catalytic domain was measured via SPR in steady-state. *Left*, fitted binding curve; *Right*, raw sensorgram data. KD shown is the mean of three independent biological repeats. Graphs shown are representatives from one experiment. (**D**) Coomassie gel of a di-ubiquitin (diUb) cleavage assay using K48-linked diUb with 50 µM WEHI-092. Representative gel of three independent biological repeats. (**E**) Coomassie gel of a ubiquitin-propargylamide (Ub-PA) suicide probe competition assay with 50 µM WEHI-092. Representative gel of three independent biological repeats. (**F**) DUB selectivity panel for WEHI-092 (50 µM) against recombinant protein performed at Ubiquigent™ (Dundee, UK). Data shown is the mean of two technical replicates which are shown as individual datapoints. (**G**) DUB IP-MS selectivity panel for WEHI-092 (50 µM) in MCF-7 cell lysates. 3X-FLAG-Ub-VS labelled DUBs were subjected to quantification by MS and % activity remaining was calculated from protein intensities relative to untreated control condition. DUBs shown have been pre-filtered to respond to Ub-VS probe pre-incubation. Data shown is the mean of five biological replicates which are shown as individual datapoints.

We next tested compound specificity using a commercial screen against a human DUB panel (see **Methods**). At 50 µM compound concentration, WEHI-092 did not affect any of 50 human DUBs with exception of USP9X (**Fig. 1F**). Compound specificity was further corroborated in cell lysates. A triple-FLAG ubiquitin-vinyl sulfone (3X-FLAG Ub-VS) suicide probe (Borodovsky *et al*, 2001), incubated with an MCF-7 cell lysate, immunoprecipitated, identified and quantified 51 human DUBs by mass-spectrometry (MS). We performed a pre-incubation of the lysate with untagged Ub-VS to reveal catalytically active DUBs in the lysate, and probe-unresponsive background levels. Out of the 51 identified DUBs, 23 responded to Ub-VS pre-incubation, suggesting that they were catalytically competent. Within these, only USP9X was affected by WEHI-092 treatment (**Fig. 1G**); the compound reduced activity levels of USP9X to the same degree as pre-incubation with unlabelled Ub-VS probe, suggesting WEHI-092 had inhibited all active USP9X in the lysate (**Fig. S1F**).

### WEHI-092 binds within the Fingers subdomain of USP9X

In the absence of mechanistic data for compound binding and specificity, we set out to understand where USP9X binds WEHI-092 and FT709. We used HDX-MS to study time dependent changes in hydrogen accessibility upon compound binding. Even at the shortest deuterium exchange timepoints (6 s), compound incubation led to protection of five overlapping peptides that included residues 1757-1762 (**Fig. 2A**). This indicated that amide protons were shielded from solvent exchange, likely due to compound interaction. These changes were more pronounced at later timepoints of the assay (**Fig. 2B**); only after 10 min deuterium exchange were changes in additional regions of USP9X detected (**Fig. S2A**). Similar results were obtained for FT709, suggesting that both scaffolds utilise the same binding site (**Fig. S2B**).

**Fig. 2.**
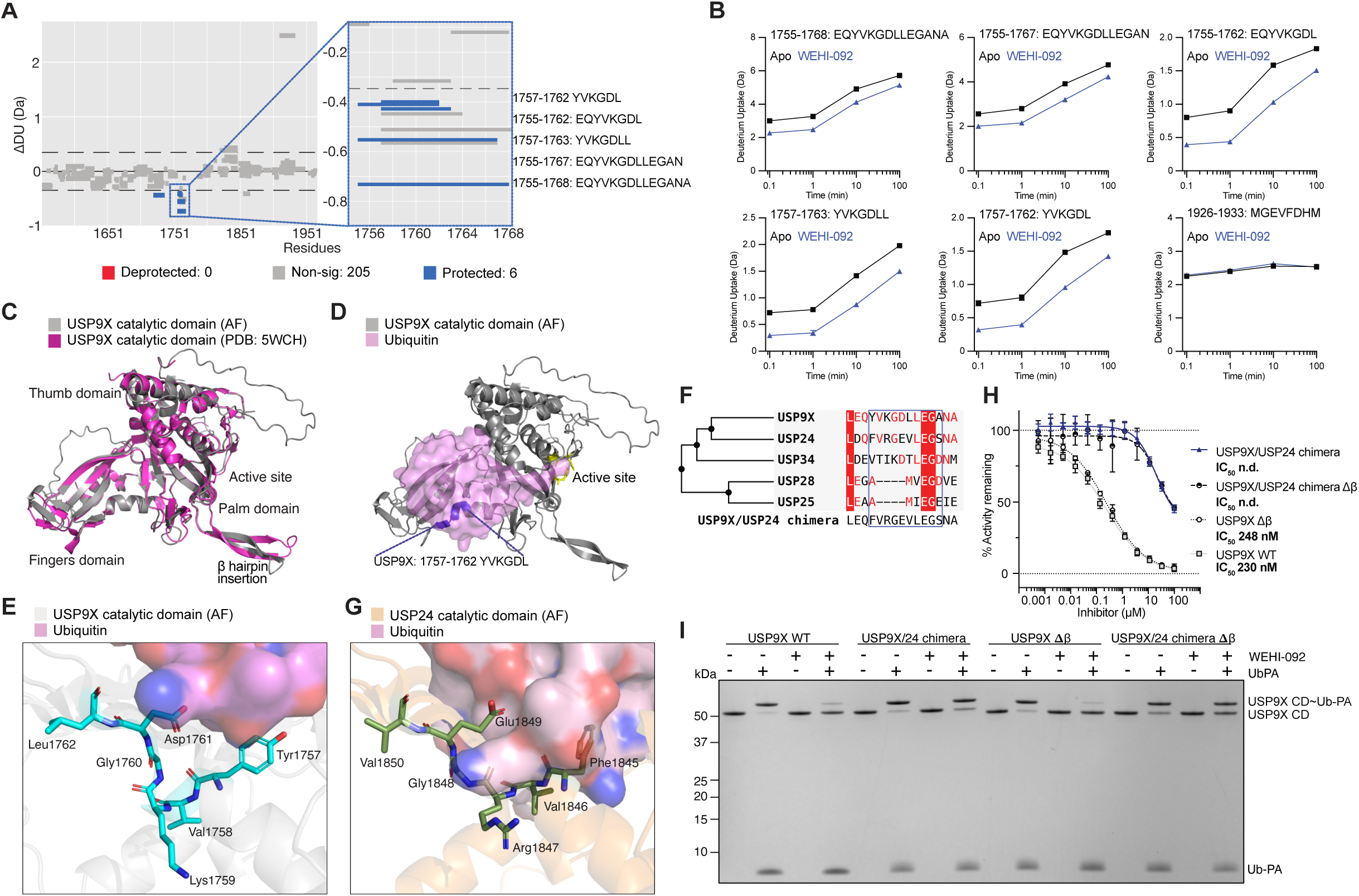
Molecular basis of WEHI-092 specificity for USP9X over other closely related USPs. **(A)** Hybrid Woods differential plot highlighting significant differences (*p* < 0.01, in blue) in hydrogen-deuterium uptake between USP9X catalytic domain apo and WEHI-092 bound protein, for 6 s timepoint. The zoomed in section highlights overlapping peptides. Dashed lines indicate the confidence limit at 99% (hybrid significance test, significance confirmed by a Welch’s t-test). Plots were generated using Deuteros software (v2.0). (**B**) Mean relative deuterium uptake plots for significant (*p* < 0.01) peptides of USP9X catalytic domain apo (black) and WEHI-092 bound (blue) from **A**, including a control peptide (ß-hairpin loop, 1926-1933, residues MGEVFDHM). Error bars, SD from n=3 experiments. (**C**) Structure of the USP9X catalytic domain (PDB: 5WCH, shown in pink) and AlphaFold3 prediction (shown superposed in grey). (**D**) Mapping of shortest peptide from **A**, YVKGDL (in blue) on the USP9X catalytic domain, with ubiquitin modelled as a pink transparent surface from an AlphaFold3 prediction. Active site residues in yellow. (**E**) Detailed view on the WEHI-092 binding site (residues YVKGDL), with blue nitrogen and red oxygen atoms, with transparent surface of modelled ubiquitin (pink carbon atoms). (**F**) Phylogenetic tree of the closest USP relatives to USP9X for the WEHI-092 binding region. (**G**) Detailed view as in **E** of the corresponding motif in the USP24 catalytic domain from an AlphaFold3 prediction. (**H**) Ub-Rho cleavage assay (as in Fig. 1B) with WEHI-092 against recombinant USP9X variants: USP9X wild-type (WT), USP9X/USP24 chimera (USP9X with 5 USP24-like mutations, see **Methods** and **F**), USP9X Δβ (USP9X β-hairpin deletion), and USP9X/USP24 chimera Δβ (USP24-like mutant with β-hairpin deletion). Data shown is the mean of three independent biological repeats with two technical replicates per experiment. IC50 values were calculated using GraphPad Prism (v10.3). n.d., not determined; error bars, SEM. (**I**) Ub-PA assay (as in Fig. 1E) using USP9X variants from **H** with and without WEHI-092. Representative gel of three independent biological repeats.

Several structures of USP9X have been reported, including a crystal structure of the isolated catalytic domain (Paudel *et al*, 2019). We modelled a ubiquitin bound catalytic domain USP9X using AlphaFold3 (Abramson *et al*, 2024) (**Fig. 2C**) and mapped the perturbed peptide motif onto this structure (**Fig. 2D**). Interestingly, the affected peptide of USP9X lies in the Fingers subdomain of the enzyme that is remote from the catalytic centre. USP catalytic domains have a characteristic ‘right-hand’-like fold, in which the ubiquitin is clasped between Fingers and Palm subdomains and cleaved within a groove provided between Palm and Thumb subdomains (Ye *et al*, 2009; Hu *et al*, 2002). All known selective USP DUB inhibitors bind residues within 12 Å on the catalytic Cys, whereas the closest distance between a residue perturbed by WEHI-092 and USP9X catalytic Cys1566 is 31 Å (**Fig. S3A, B**). Moreover, the perturbed peptide is located at the edge of the Fingers subdomain’s β-sheet, without forming a notable pocket or groove, yet some residues are in hydrogen binding distance with ubiquitin (**Fig. 2E**). Additional peptides within the Fingers subdomain appear to be deprotected at later timepoints potentially suggesting long-range compound induced conformational changes (**Fig. S2A).** Biochemical experiments (**Fig. 1**) show that compound binding affects ubiquitin interactions, which would be consistent with WEHI-092 binding to the Fingers domain as shown through an AlphaFold3 prediction with ubiquitin bound (**Fig. 2E**).

We used phylogenetic analyses and mutagenesis to corroborate the compound binding site. USP9X has a close human paralogue, USP24, a less-well studied enzyme (Wang *et al*, 2020; Rossio *et al*, 2024) with a highly similar overall domain architecture, in which catalytic domains are embedded in expansive helical repeat scaffolds (**Fig. S2E**). The catalytic domain shares a sequence identity of 47% (sequence similarity 61%) with similar overall fold, also featuring the characteristic extended β-hairpin. USP24 was included in DUB specificity panels tested against WEHI-092, but was not inhibited by the compound neither in the commercial panel nor in cell lysates (**Fig. 1F, G**). The perturbed peptide within USP9X (aa: YVKGDL) differs in sequence, but not in length in USP24 (aa: FVRGEV) (**Fig. 2F, G**). We hence created a chimera, in which USP9X residues were replaced with USP24 residues in a 11-mer sequence, in a wild-type (WT) catalytic domain, and in a catalytic domain in which the extended β-hairpin, a unique feature of USP9X and USP24, was removed (USP9X Δβ). Individual mutations slightly reduced USP9X activity in a Ub-Rho assay as has been observed before (Paudel *et al*, 2019), yet each variant remained robustly active (**Fig. S2D**). Importantly, while wild-type USP9X and USP9X Δβ were inhibited by WEHI-092, the chimera mutant catalytic domains remained fully active at concentrations up to 10 µM, and partially active even at the highest compound concentrations (**Fig. 2H**). At 50 µM compound concentration in a Ub-PA suicide probe assay, WT or USP9X Δβ catalytic domains are unable to be modified by the probe, whereas chimera mutants still bind and are modified by Ub-PA probes (**Fig. 2I**). Pinpointing a short stretch of amino acids to be involved in compound binding reveals a new USP9X compound binding site in the Fingers domain distinct from other selective USP DUB inhibitors (**Fig. S3A, B)**.

### WEHI-092 is cytotoxic in a small subset of cancer cell lines

Next, we tested the inhibitory effects of WEHI-092 on the proliferation and survival of a panel of cancer cell lines. To ensure compound activity in cells, we tested a previously established proximal biomarker, the centrosomal protein CEP55 (Wang *et al*, 2017). CEP55 was destabilised in a dose-dependent fashion by FT709 as reported previously (Clancy *et al*, 2021) and also by WEHI-092, with an apparent IC50 (based on Western blotting) of 1.3 µM (**Fig. S4A**).

We assayed cell lines that included a broad range of cancer types which were all found to express USP9X to varying degrees (**Fig. S4B**). To understand the impact of USP9X inhibition on cancer cell lines, we initially performed a series of cellular assays by treating cells with WEHI-092 (**Fig. 3A**). These included (i) short term (3-day) cell killing/toxicity studies and (ii) short term (3-day) cell growth/proliferation assessment, using the cancer cell line panel provided by the National Cancer Institute (NCI) (Shoemaker, 2006) in which we tested WEHI-092 at 10 µM (a concentration at which clear target regulation was observed consistently, see below), against 57 human cancer cell lines across nine different cancer types (**Fig. 3B**). A 10-fold less potent compound, WEHI-680, was tested for comparison (**Fig. S4C**). The results from the NCI panel were confirmed and expanded to additional cancer cell lines as well as non-cancerous cell lines (HUVECs and human dermal fibroblasts, HDFs), by performing in-house dose-escalation studies using IncuCyte live-cell imaging analysis, assessing cell death (**Fig. 3C**) and cell growth (**Fig. 3D**). These broad analyses of compound effects performed across multiple cell types revealed several interesting results. Firstly, WEHI-092 is non-lethal and non-toxic to the non-cancerous human cell lines studied (HUVECs and HDFs), a finding that was distinct from commonly used non-specific DUB inhibitors WP1130 and EOAI3402143, which were cytotoxic to HDFs (**Fig. S4E**).

**Fig. 3.**
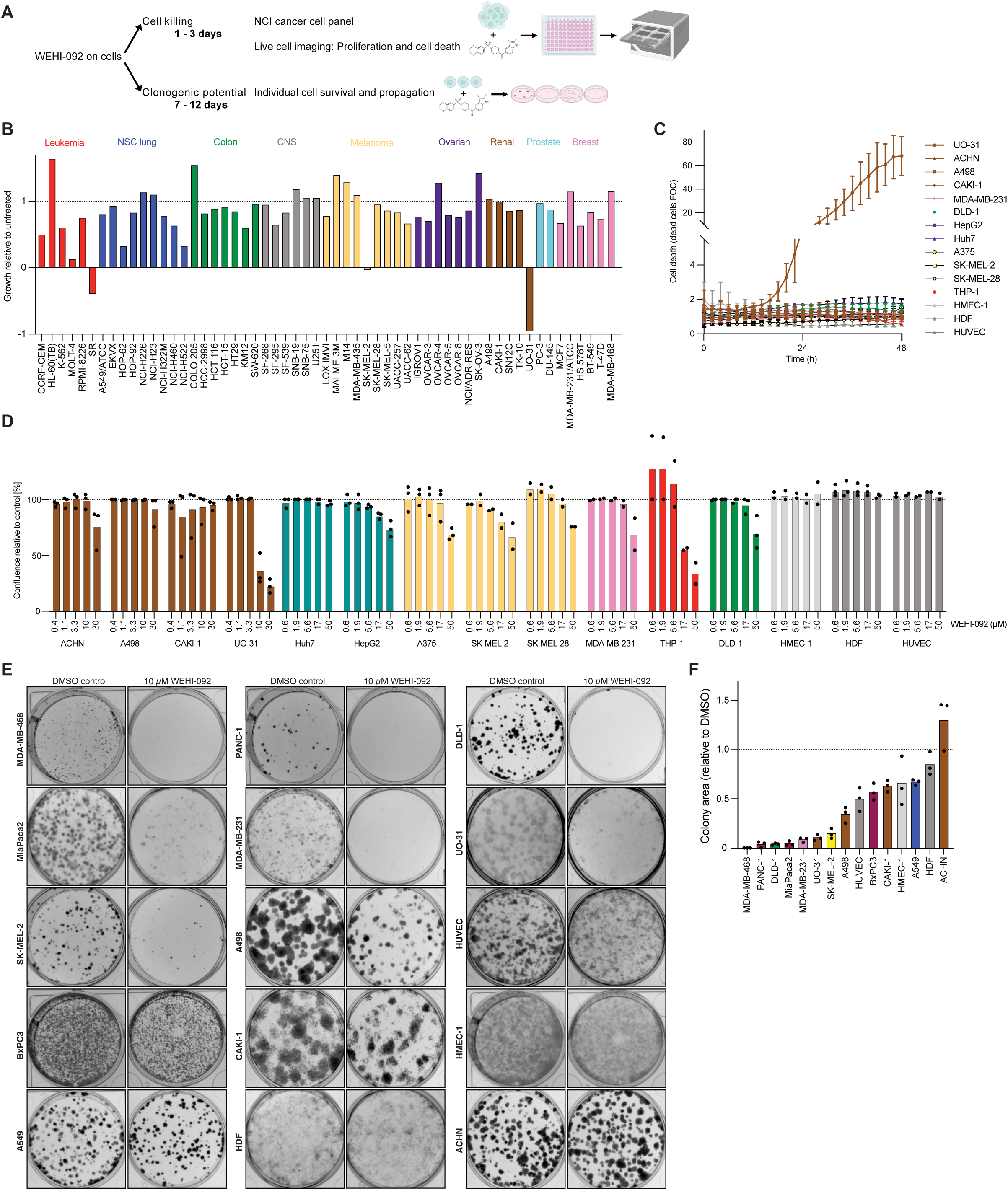
Treatment with WEHI-092 induces cell line specific cell killing and clonogenic potential phenotypes. (A) Schematic of short-term cell killing and long-term clonogenic potential experiments. (B) NCI-60 cancer cell panel of 57 cancer cell lines treated with WEHI-092 (10 µM). Data shown is the normalised cell growth at 3 days post treatment relative to untreated control and relative to day 0 for each individual cell line. Negative values indicate cell death at 3 days post treatment. Cell lines are colour-coded depending on their cancer type. NCS - non-small cell. CNS - central nervous system. Data shown is the mean of two technical replicates. (**C**) Incucyte live cell imaging cell death data for cell lines treated with WEHI-092, used at 10 µM for UO-31, A498, ACHN, CAKI-1, HMEC-1, HDF cells and at 17 µM for the remaining cell lines. Cell death was measured by propidium iodide uptake. Data shown is the mean of two or three independent biological repeats with two technical replicates. Error bars, SEM. FOC - fold over control. HDF - Human dermal fibroblasts (primary). (**D**) Incucyte live cell imaging proliferation data for cell lines treated with WEHI-092. Data shown is the mean of two to three independent biological repeats with two technical replicates at 72 h post treatment where each experiment is shown as a single datapoint. Data was normalised to the untreated condition. (**E**) Clonogenic potential assay for cell lines treated with WEHI-092. Representative images from three independent biological repeats with three technical replicates. (**F**) Quantification of clonogenic potential assays as shown in **E**. Data shown is the mean of three independent biological repeats with three technical replicates relative to the untreated condition (dashed line) for each cell line where each separate experiment is shown as a single datapoint. Bars are colour-coded as in **B**.

Secondly, USP9X inhibition showed little to no effect in most cell lines tested in 3-day assays. With few exceptions, cell growth was only mildly impaired or not affected. However, in five cell lines across three cancer types, WEHI-092 caused more than 50% cell growth inhibition, including stalling cell growth completely in MOLT-4 and SK-MEL-2 cells (**Fig. 3B**). Thirdly, WEHI-092 caused cell death of the lymphoma line SR, and of the papillary renal cell carcinoma (pRCC) UO-31 cell line (Sinha *et al*, 2017), with the latter demonstrating unique sensitivity to WEHI-092. Consistently, UO-31 cells were the only cell line in the panel that was killed by WEHI-092 in IncuCyte assays (**Fig. 3C**). Other RCC cell lines, including clear cell renal cell carcinoma (ccRCC) lines but also a second pRCC line, ACHN, showed limited response to WEHI-092 treatment (**Fig. 3C**). Notably, UO-31 cells were highly sensitive to WEHI-092 but less sensitive to FT709 (**Fig. S4F**), with the slightly better *in vitro* activity of FT709 not translating to improved cellular activity (**Fig. S4A**).

Next, we assessed (iii) longer-term (7-14 days) effects of WEHI-092 on the clonogenic potential of cancer cells, either at a single compound concentration (10 µM, **Fig. 3E, F**) or in a dose-escalation study, the latter in comparison with WEHI-680 (**Fig. S4D**). As previously observed in the short-term assays, SK-MEL-2 and UO-31 cells were sensitive to WEHI-092 treatment (**Fig. 3E**). However, we also observed a strong decrease in proliferative capacity for the colorectal cancer cell line DLD-1, the pancreatic cancer cell lines MiaPaca2 and PANC-1 as well as breast cancer cell lines MDA-MB-231 and MDA-MB-468 (**Fig. 3E, F**). Other cancer cell lines tested, and also non-cancer cells (HUVEC and HDFs) showed limited or no effects in clonogenic potential suggesting cell type specific effects of USP9X inhibition. This data suggested that extended treatment time exacerbates cell growth defects in cultured cell lines. It is also worth noting that we have not observed an increase in cell proliferation, in any cell line studied.

### USP9X and USP7 have a non-overlapping set of substrates

The disparate rates of survival and cellular proliferation between cancer cell lines in response to WEHI-092 was unexpected and warranted a deeper investigation of USP9X and its regulatory roles, using a specific DUB inhibitor. We hence turned to comparative ubiquitinomics and proteomics studies, to study the effects of DUB inhibitors across cell lines. First, we sought to test whether different DUBs regulate the same set of ubiquitination sites within a given cell line, or whether they indeed exhibit substrate specificity, as is widely assumed in the field. To date, this has only been explored indirectly, at the transcriptomic levels using RNAseq which suggested a non-overlapping set of substrates for USP7, USP14, and USP30 (Doherty *et al*, 2022).

Steger et al. recently performed comprehensive proteomic and ubiquitinomic studies of several chemically distinct USP7 inhibitors, including the highly specific and potent FT671 (Turnbull *et al*, 2017), in a colorectal adenocarcinoma cell line (HCT116) (Steger *et al*, 2021). Using FT671 in a similar colorectal carcinoma cell line (DLD-1) with identical concentration (10 µM), time points (30 min, 360 min), and liquid-chromatography (LC)-MS acquisition, we detected 23,568 ubiquitinated GlyGly (K-GG) peptides that were reported by Steger et al. (**Fig. 4A**). Filtering for K-GG peptides that were significantly increased during FT671 treatment in Steger et al. revealed good quantitative reproducibility between the two studies (**Fig. S5A, B**), which was also apparent at the protein level for significantly reduced proteins following FT671 treatment (**Fig. S5C**).

**Fig. 4.**
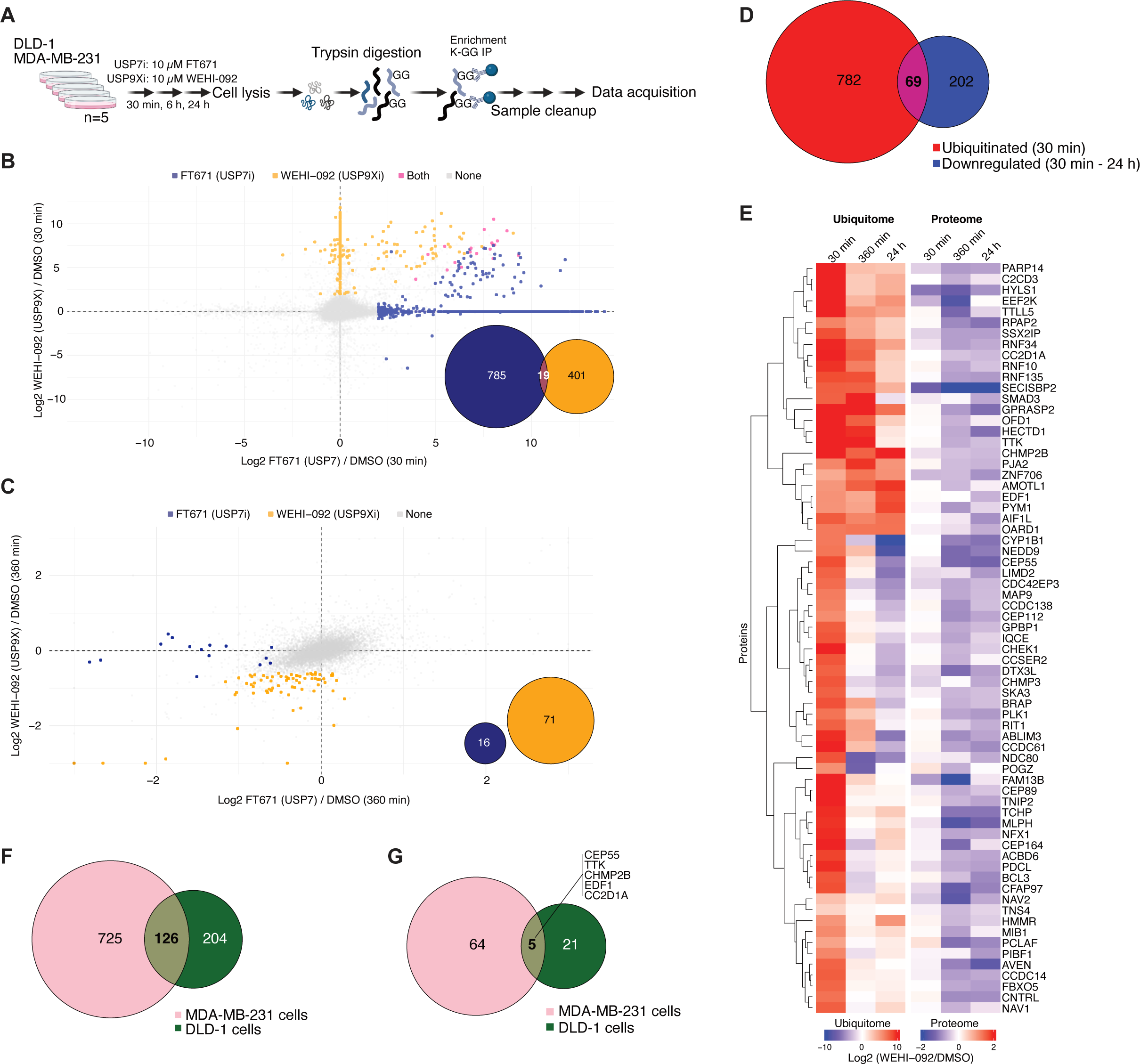
Time resolved ubiquitome profiling reveals high confidence USP9X substrates in a cell line dependent manner. **(A)** Schematic overview of the ubiquitome profiling experiments. (**B**) USP9X vs. USP7 inhibition in DLD-1 cells treated with FT671 (USP7 inhibitor) or WEHI-092 at 10 µM for 30 min. Each datapoint shows the mean log2 fold change for USP7i or USP9Xi treatment vs. control condition across five technical repeats. Coloured datapoints show K-GG peptides with a fold change of log2 > 2 (FDR < 0.05) over untreated control in blue for USP7i, in yellow for USP9Xi and in pink in both treatments. Empty values of non-detected peptides in one or the other condition were filled with zero values and are clustered on the x-and y-axis respectively. The Venn diagram shows the number of K-GG peptides with a fold change of log2 > 2 (FDR < 0.05) over untreated control for the respective conditions. (**C**) USP9X vs. USP7 inhibition in DLD-1 cells treated with FT671 or WEHI-092 at 10 µM for 360 min. Each datapoint shows the mean log2 fold change upon USP7i or USP9Xi treatment vs. control condition across 5 technical repeats. Coloured datapoints show proteins with a fold change of log2 < -0.585 (FDR < 0.05) over untreated control in blue for USP7i, in yellow for USP9Xi. Non-detected proteins in one or the other condition were filled with zero values and missing values were imputed using a mixed BPCA and Min method. Axis limits were set to -3 / 3 and datapoints outside of the range were plotted on the axis limit. The Venn diagram shows the number of downregulated proteins with a fold change of log2 < -0.585 (FDR < 0.05) over untreated control for the respective conditions. (**D**) Venn diagram showing the overlap of proteins with enhanced ubiquitination at 30 min (log2 > 2 fold, FDR < 0.05 over untreated control) and decreased abundance at either 30 min, 360 min or 24 h WEHI-092 treatment (10 µM) (log2 < -0.585, FDR < 0.05 over untreated control) in MDA-MB-231 cells. (**E**) Time resolved profiling of high confidence USP9X substrates in MDA-MB-231 cells. Heatmap colours indicate fold change in protein ubiquitination (left) and protein expression (right) of proteins that showed significant induction (fold change log2 > 2, FDR < 0.05 over untreated control) of at least one ubiquitination site at 30 min of WEHI-092 treatment (10 µM) and that were significantly downregulated (fold change log2 < -0.585, FDR < 0.05 over untreated control) at 30 min / 360 min / 24 h. The data was averaged across five technical replicates and the data was matched based on gene ID level. Hierarchical clustering was performed on proteins (rows) with Euclidean distance as the similarity metric. (**F**) Comparison of ubiquitinated proteins (log2 > 2-fold increase over untreated control, FDR < 0.05) after 30 min of WEHI-092 treatment between MDA-MB-231 and DLD-1 cells. (**G**) Comparison of high confidence USP9X substrates in MDA-MB-231 cells and DLD-1 cells. High confidence USP9X substrates were identified in MDA-MB-231 cells from **E** and in DLD-1 cells (**Fig. S5D**) and are defined as upregulated ubiquitination (fold change log2 > 2, FDR < 0.05 over untreated control) and depleted protein levels (fold change log2 < -0.585, FDR < 0.05 over untreated control).

Next, we treated DLD-1 cells with the USP9X inhibitor WEHI-092 at 10 µM. A distinct set of K-GG peptides were significantly increased with USP9X inhibition (using WEHI-092) as compared to USP7 inhibition (using FT671) at 30 min of treatment (**Fig. 4B**); 804 K-GG-modified peptides were enriched upon USP7 inhibitor treatment, 420 K-GG-peptides upon WEHI-092 treatment, and only 19 K-GG-peptides increased with either treatment (**Fig. 4B**). Moreover, there was no overlap in proteins that were significantly reduced between USP7 inhibitor treatment and USP9X inhibitor treatment (**Fig. 4C**). These results revealed the significant substrate specificity of two distinct, individually highly abundant USP DUBs in cells at the protein level.

### Defining high confidence substrates of USP9X by ubiquitinomics and proteomics

Steger et al. reported that USP7 inhibition led to increased ubiquitination within 15-30 min, which corresponded to reduced abundance of the ubiquitinated proteins after 6 h of USP7 inhibition (Steger *et al*, 2021). In order to identify high confidence substrates (increased ubiquitination and decreased protein abundance following DUB inhibitor treatment) of USP7 and USP9X in DLD-1 cells, proteins were filtered for possessing K-GG peptides that were significantly increased at 30 min (log2 > 2 fold change, false discovery rate (FDR) < 0.05) and the total protein pool reduced after 6 h of DUB inhibition (log2 < -0.3, FDR < 0.05) (**Fig. S5D, E**). With either inhibitor (and as previously observed (Steger *et al*, 2021)), the number of ubiquitinated proteins exceeded the number of subsequently degraded proteins, by at least ∼6-fold. This observation may indicate either non-degradative (e.g. signalling) ubiquitination events, or ubiquitination events that are not yet functional to induce proteasomal degradation. 26 proteins from DLD-1 cells were ubiquitinated at short time points and degraded at longer time points meeting the criteria as high confidence USP9X substrates (**Fig. S5D**).

We expanded our analysis of USP9X inhibition by profiling the ubiquitin landscape of another cancer cell line to understand the consistency of high confidence USP9X substrates between two distinct cancer cell lines. The breast cancer cell line MDA-MB-231 was treated identically (10 µM WEHI-092, measured at 30 min, 6 h, 24 h). 30 min USP9X inhibition increased the abundance of 1323 K-GG peptides on 851 proteins, while across all time points, a total of 271 proteins were significantly decreased (**Fig. 4D**). 69 proteins exhibited increased ubiquitination at 30 min and decreased protein levels (following 30 min, 6 h and/or 24 h inhibition) upon WEHI-092 inhibition, indicating high-confidence USP9X substrates (**Fig. 4E**). Among these targets are several previously reported USP9X substrates such as MIB1 (Izrailit *et al*, 2017), TTK (Chen *et al*, 2018), SMAD3 (Dupont *et al*, 2009), RIT1 (Riley *et al*, 2024), and CEP55 (Wang *et al*, 2017; Clancy *et al*, 2021) that were matched to corresponding biological processes through pathway enrichment analysis (**Fig. S5F, G**).

Interestingly, comparing substrates between different cell lines revealed only a partial overlap of ubiquitinated proteins at 30 min USP9X inhibition (**Fig. 4F**) and only 5 proteins (CEP55, TTK, CHMP2B, EDF1, CC2D1A) that were regulated in both, MDA-MB-231 and DLD-1 cells (**Fig. 4G**). Indeed, of the 69 high confidence USP9X substrates found in MDA-MB-231 cells, 58 were detected in DLD-1 cells at the peptide level, but were seemingly not significantly altered by USP9X inhibitors during the experiment.

### Expansion of proteomic studies to eight cell lines reveals a core set of commonly depleted USP9X targets

While sets of high confidence USP9X substrates emerged in two cell lines treated with WEHI-092, the disparity of putative substrates was unsettling. To further corroborate findings, we first validated target lists against published data generated with USP9X inhibitor FT709. Treatment of HCT116 cells with FT709 significantly reduced CEP55, MKRN2, TTK, CEP131, PCM1 and other proteins (Clancy *et al*, 2021). We extended these proteomics analyses to MDA-MB-231 cells, which were treated with FT709 or WEHI-092, for 24 h. Notably, also treatment with WEHI-092 led to reduced protein levels of CEP55, MKRN2, CEP131 and PCM1, producing highly correlated datasets overall (**Fig. 5A**).

**Fig. 5.**
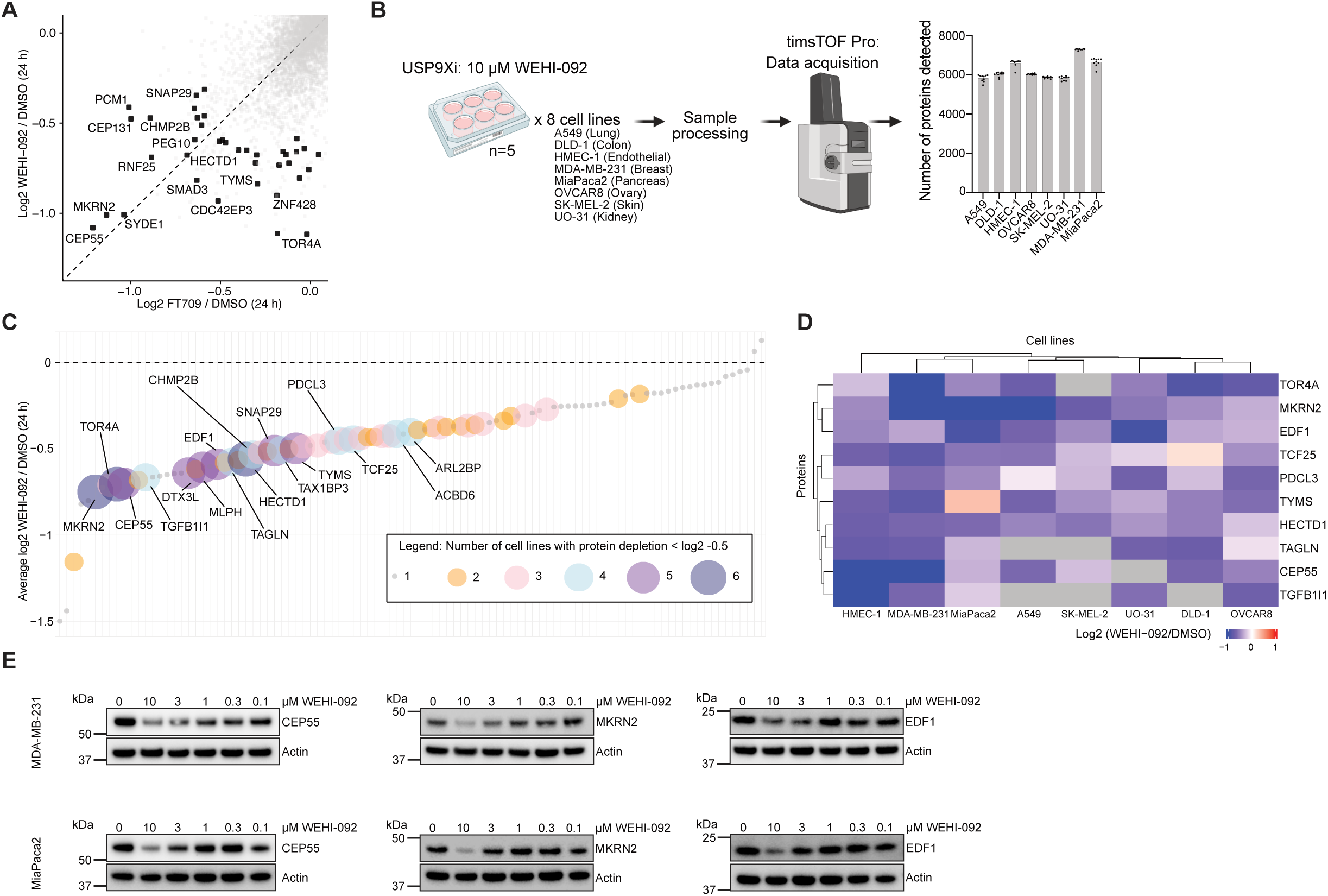
Global proteomics studies identify commonly regulated proteins across cell lines upon USP9X inhibition. **(A)** Comparison of depleted proteins (log2 < -0.585-fold change over untreated control, p.adj < 0.05) in MDA-MB-231 cells upon USP9X inhibitor treatment with either FT709 or WEHI-092 (10 µM, 24 h). Each datapoint represents the mean of five technical replicates. The dashed line at y=x was included for visualisation purposes. (**B**) Schematic workflow of global proteomics experiments in eight cell lines. The bar graph on the right shows the number of detected proteins per cell line (mean). Each datapoint represents one of 5 technical repeats per condition (with and without WEHI-092). (**C**) Identification of commonly regulated proteins upon WEHI-092 treatment (10 µM, 24 h) across the eight cell lines. Plotted is the average log2 fold change over untreated control across all eight cell lines, with n=5 per cell line per condition. The size of each dot indicates in how many cell lines the respective protein was depleted by log2 < -0.5-fold change WEHI-092 treated vs. untreated control treatment. The data was pre-filtered for proteins that are depleted by log2 < -0.585 (p.adj. < 0.05) over untreated control in at least one out of the eight cell lines. Proteins that are depleted in at least four / eight cell lines are labelled. The y-axis limit was set to -1.5 and one datapoint outside of this was plotted at y = -1.5 (**D**) Heatmap visualisation of the top ten commonly regulated proteins. Data from **C** was filtered for the top ten proteins with a significant (p.adj < 0.05) protein level decrease by log2 < -0.585 over untreated control in at least three out of the eight cell lines. Hierarchical clustering was performed on proteins (rows) and cell lines (columns) using Euclidean distance as the similarity metric. Heatmap colours indicate fold changes in protein expression, whereby grey indicates that the protein was not detected. (**E**) Western Blot validation of a subset of commonly regulated USP9X substrates from **C** and **D**. Treatments were performed for 24 h. Representative blots from at least two independent biological repeats are shown.

We next expanded the global proteomic studies to six additional cancer cell lines, covering different phenotypes seen in the NCI panel, including cell lines with no growth defect, growth reduction, and including UO-31 cells that were highly sensitive to WEHI-092 treatment (**Fig. 5B**, compare **Fig. 3B, C**). This set of proteomic experiments were performed on Bruker timsTOF MS instrumentation (see **Methods**), which achieved proteome coverage from 5833 (UO-31) to 7310 proteins (MDA-MB-231) (**Fig. 5B**). The re-run of MDA-MB-231 cells overlapped in 6828 out of 7310 proteins compared with the first analysis (**Fig. 4D**, performed on Thermo Astral MS instrument), in which 9232 proteins were observed. All cell lines were treated with 10 µM of WEHI-092 or DMSO (control) for 24 h, differential proteins extracted individually for each cell line, and cross-compared across cell lines.

Filtering for significantly reduced proteins across each cell line indicated that 17 core proteins are decreased in at least 4 out of 8 cell lines tested (**Fig. 5C**). The top ten regulated proteins by USP9X inhibition include some previously validated targets including CEP55 and MKRN2, and further proteomic hits (EDF1, TCF25) previously identified (Clancy *et al*, 2021), while TOR4A, PDCL3, TYMS, HECTD1, TAGLN and TGFB1I1 were consistently depleted across cell lines but have not previously been associated with USP9X (**Fig. 5D**). The reduction in CEP55, MKRN2 and a new target, EDF1 following USP9X inhibition in multiple cell lines were confirmed by Western blotting (**Fig. 5E, Fig. S6B**). These studies also showed that USP9X levels were not affected by compound treatment, and that CEP55 depletion was rescued by proteasome inhibition whenever assessed (**Fig. S6A**), consistent with ubiquitin-dependent turnover and direct, rather than indirect (transcriptional/translational) regulation. In addition to decreases in protein levels, several of the top ten downregulated proteins across all cell lines had corresponding ubiquitinated peptides in the ubiquitinomics studies including CEP55, EDF1, MKRN2, HECTD1 (**Fig. 4D, E, Fig. S5D**). The identification of specific proteins that were consistently ubiquitinated and depleted between cell lines, revealed a valuable set of biomarkers for USP9X inhibition.

### Global analysis of USP9Xi regulated proteins reveals cell line specific patterns with functional clusters

Analysis of proteomic datasets for individual cell lines revealed a relatively small number of proteins (between 5 to 30) that changed significantly upon USP9X inhibition at the chosen time point. To identify global patterns of proteome changes by USP9X inhibition using pathway analysis, we combined the results from all cell lines, analysing proteins that were significantly downregulated in at least one of the cell lines. This generated a list of 99 proteins, which were then visualised in context of their cell line origin (**Fig. 6A**). Strikingly, while there was a set of overlapping proteins (which included proteins discussed above and in **Fig. 5**), the majority of proteome changes occurred in a cell-line specific fashion (**Fig. 6A**). The observed depletion upon USP9X inhibition could not be explained by abnormal expression patterns of targets across the tested cell line (**Fig. S7A**). This data was very surprising, as it suggested disparate cell type-specific roles of USP9X. Such cell-type and context specific effects, may potentially explain the fact that many of the previously suggested targets of USP9X (such as MCL-1 (Schwickart *et al*, 2010), ERG (Wang *et al*, 2014), Ets-1 (Potu *et al*, 2017), YAP1 (Li *et al*, 2018; Biber *et al*, 2024), NUAK1 (Fritz *et al*, 2020), and Itch (Mouchantaf *et al*, 2006)) were not observed as significantly changing proteins in any of the cell lines here tested (see **Discussion**). We next undertook enrichment analysis of the proteomic changes across different cell lines to understand the pathways and cellular processes affected by USP9X wherein individual proteins were systematically categorised by gene ontology (GO) terms and subsequently aggregated by biological pathways into clusters (Zhou *et al*, 2019). Stringent filtering criteria were applied (protein level reduction > 50%, p.adj < 0.05) to identify enriched GO-terms associated with the most significant hits across cell lines. In this enriched ontology map, nodes representing similar biological processes and pathways were identified and clustered (**Fig. 6B**).

**Fig. 6.**
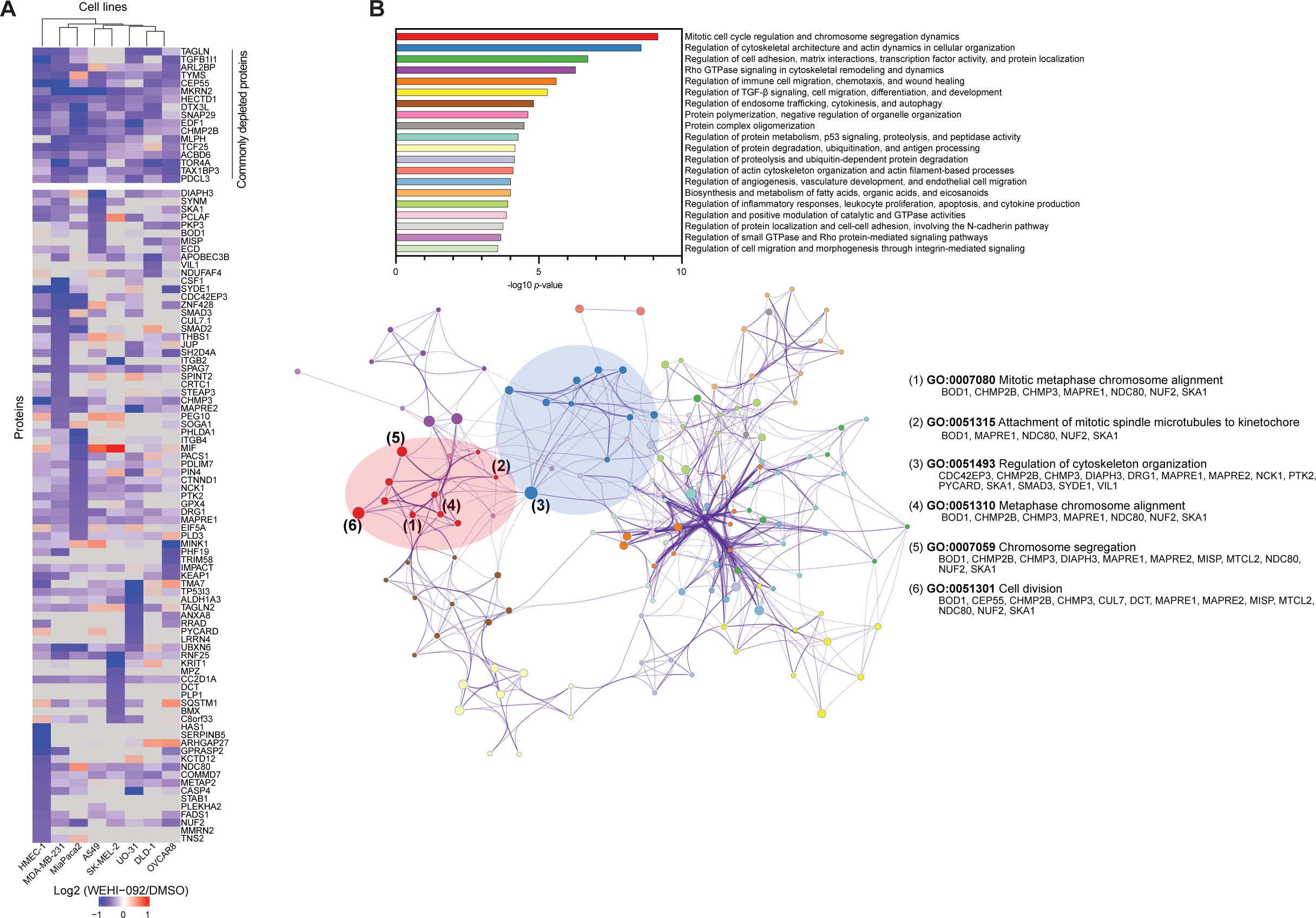
Global proteomics across eight different cell lines reveals cell line specific regulation of proteins upon WEHI-092 treatment. **(A)** Heatmap visualisation of proteins with altered abundance upon WEHI-092 treatment in eight cell lines. The data was pre-filtered for proteins that are depleted by log2 < -0.585 (p.adj. < 0.05) over untreated control in at least one out of the eight cell lines. Five technical replicates were included in the experiment per cell line. Hierarchical clustering was performed on cell lines (columns) using Euclidean distance as the similarity metric. Heatmap colours indicate log2 fold changes in protein expression, whereby grey indicates that the protein was not detected. 17 proteins in the top part are commonly depleted (see Fig. 5). (**B**) Pathway and process enrichment analysis of proteomic dataset from (a) using metascape.org (Zhou *et al*, 2019). Similar biological processes and pathways were clustered algorithmically, colour-coded for differentiation, and summarised. Clusters were quantitatively ranked in a bar plot (top) and visualised as a network (bottom) where each node corresponds to a specific gene ontology term, with node diameters proportional to the number of contributing proteins. Nodes representing similar biological processes and pathways from the same cluster are shown in the same colour used in the bar plot. Within the network, nodes from within the top two clusters are highlighted through a colour backdrop. The top six most significant individual gene ontology terms are numbered and labelled with their respective gene ontology term and with the protein names from our dataset falling into the term.

Analysis of all 99 proteins across cell lines, unveiled distinct biological pathways with which USP9X has previously been associated, including, TGFβ-signalling (Dupont *et al*, 2009; Jie *et al*, 2021; Yang *et al*, 2022), ubiquitin-dependent catabolism (Xie *et al*, 2013; Mouchantaf *et al*, 2006), cytoskeleton architecture (Homan *et al*, 2014) and endosomal trafficking (Savio *et al*, 2016). The most prominent cluster of regulated proteins upon USP9X inhibition related to GO terms, concerned mitotic cell cycle regulation and chromosome segregation (**Fig. 6B**) consistent with the reported role of USP9X in cell cycle regulation and mitosis (Skowyra *et al*, 2018; Dietachmayr *et al*, 2020; Engel *et al*, 2016; Kodani *et al*, 2019; McGarry *et al*, 2016; Li *et al*, 2017). Specifically, in addition to the previously reported CEP55 which is depleted across all cell lines, we find multiple centrosomal/kinetochore proteins including Biorientation Of chromosome in cell Division protein 1 (BOD1), Spindle and Kinetochore Associated complex subunit 1 (SKA1) and kinetochore complex component NDC80, to be USP9X targets, albeit in different cell lines (**Fig. 6B**).

### USP9X inhibition arrests cells in metaphase

To functionally validate our global proteomic findings, we selected three cell lines from our panels in **Fig. 3** to study the effects of WEHI-092 in mitosis using lattice light sheet imaging. Two out of the three selected cell lines (MDA-MB-231 and MiaPaca2) showed sensitivity to compound treatment in clonogenic potential assays whereas HMEC-1 cells were largely unresponsive to WEHI-092. We analysed cell division by following individual cells over a 24 h window by imaging tubulin and DNA in cells every 10 min and tracked cell divisions with and without WEHI-092 treatment (**Fig. S8A-D**). MDA-MB-231 and MiaPaca2 cells showed significant defects in cell division following treatment with WEHI-092 (**Fig. 7A, B, D**). While the chromosomes condense and the spindle apparatus forms successfully with moderately delayed kinetics, there was no progression beyond metaphase in treated cells (**Fig. 7A, B**, **Videos S1-4**). In contrast, HMEC-1 cells which were unresponsive to WEHI-092 treatment in growth or survival assays, proceeded through mitosis normally (**Fig. 7C, D**). Notably, metaphase arrest did not trigger apoptosis in MDA-MB-231 and MiaPaca2 cells, as assessed by AnnexinV uptake (**Fig. 7E**) suggesting that USP9X inhibition induces a non-lethal mitotic blockade in susceptible cell lines likely explaining effects on clonogenic potential, and growth arrest in cancer cell lines (**Fig. 3**). Together these analyses demonstrate how meta-analysis of proteomic changes induced by a specific DUB inhibitor across various cell lines, can provide significant insights into global DUB function.

**Fig. 7.**
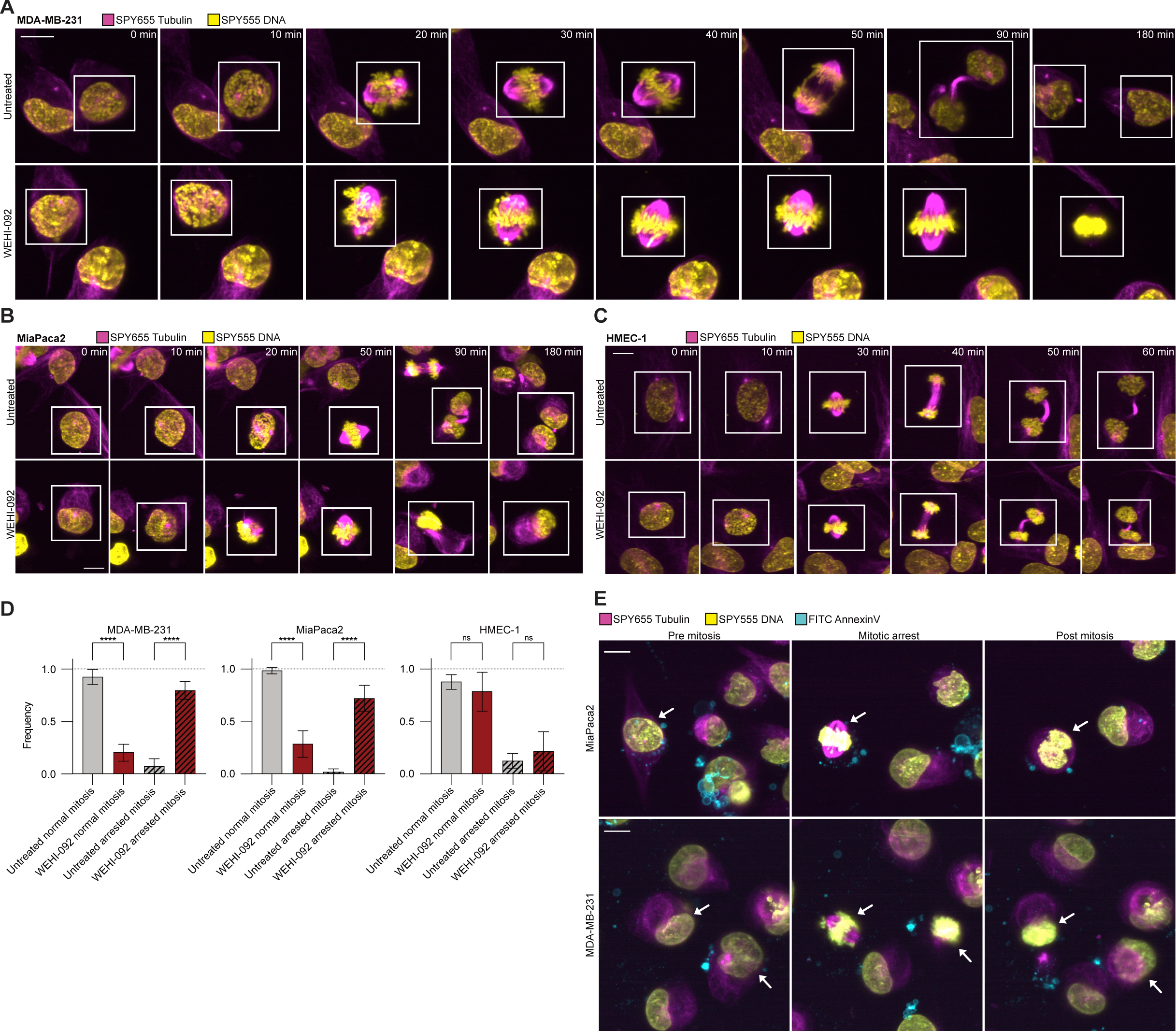
Inhibition of USP9X causes arrested mitosis in cancer cells. **(A)** Lattice light sheet imaging of MDA-MB-231 cells upon WEHI-092 treatment. Cells were treated with 15 µM WEHI-092 for 24 h before imaging commenced. Times indicated refer to initiation of mitosis for the cells highlighted in the white boxes. DNA is shown in yellow, and tubulin is shown in magenta. Images were taken every 10 min and are a representative of three independent biological repeats with a total of 29 divisions (untreated) and 25 divisions (WEHI-092) respectively. Scale bar: 10 µm. (**B**) Lattice light sheet imaging of MiaPaca2 cells upon WEHI-092 treatment. Conditions and colour channels are similar to **A**. Images were taken every 10 min and are a representative of three independent biological repeats with a total of 92 divisions (untreated) and 39 divisions (WEHI-092). Scale bar: 10 µm. (**C**) Lattice light sheet imaging of HMEC-1 cells upon WEHI-092 treatment, also as in **A**. Images were taken every 10 min and are a representative of three independent biological repeats with a total of 30 divisions (untreated) and 34 divisions (WEHI-092). Scale bar: 10 µm. (**D**) Quantification of mitosis phenotypes in MDA-MB-231, MiaPaca2 and HMEC-1 cells. Cell divisions were categorised into normal and arrested mitosis based on the phenotypes observed in **A-C**. In the bar graphs, the mean across three independent biological repeats is shown with a total of 29 divisions (untreated) and 25 divisions (WEHI-092) for MDA-MB-231 cells; 92 divisions (untreated) and 39 divisions (WEHI-092) for MiaPaca2 cells; 30 divisions (untreated) and 34 divisions (WEHI-092) for HMEC-1 cells. Error bars, SD. One-way ANOVA with post-hoc multiple comparison Tukey test was performed for significance testing. Significance level **** *p* < 0.0001, ns - not significant. (**E**) Cells with arrested mitosis phenotype are not staining positive for AnnexinV. MiaPaca2 and MDA-MB-231 cells were treated as described in **A** and **B**. DNA is shown in yellow, tubulin is shown in magenta, and AnnexinV is shown in cyan. Scale bar: 10 µm. Representative images of three independent experiments. Arrows indicate the same cell before, during and post mitosis.

## Discussion

We here identify a new chemical class of USP9X inhibitors with a distinct chemical piperazine core (Clancy *et al*, 2021) that is more chemically tractable as compared to the pyrrolo[3,4-*c*]pyrrole core in FT709, with similar *in vitro* activity, cellular efficacy, and, most importantly, remarkable USP9X specificity *in vitro* and in cells.

We explain specificity of WEHI-092 and FT709 through HDX-MS experiments, which reveal a single 6 amino acid region, that located surprisingly in the Fingers subdomain of USP9X. This 6 amino acid motif is unique to USP9X (Ye *et al*, 2009) among USP family members and differs even in its closest human paralogue, USP24. A USP9X chimera with five USP24-mimicking point mutations, was no longer able to bind WEHI-092 or FT709. Hence, we have pinpointed the binding site for USP9X specific inhibition, which remarkably is located away from the catalytic centre and represents a previously unrecognised compound binding pocket in the Fingers subdomain. Despite significant effort, we were unable to obtain a co-crystal structure, and it remains to be established how WEHI-092 precisely binds to USP9X. Yet again, these findings demonstrate that USP DUBs are remarkable for the many ways that the catalytic domain can accommodate chemically distinct scaffolds and be inhibited (Kazi *et al*, 2025).

A new USP9X inhibitor scaffold presented an opportunity to clarify the role of USP9X in cancer cell biology. Despite more than 130 papers on this topic, the USP9X field was conflicted as to whether USP9X is a viable cancer target (Dewson *et al*, 2023). By using WEHI-092 across a large panel of cancer cell lines combined with performing deep ubiquitinomic and proteomic analyses, we explain some of the discrepancies in prior literature and consolidate the literature by revealing key pathways regulated across cell lines and cancer settings. We find that USP9X inhibition affects a unique set of proteins as compared to USP7 inhibition, which clearly demonstrates that these two abundant and ubiquitous DUBs have distinct sets of protein substrates. USP9X inhibition increases the abundance of over 1000 ubiquitination sites in > 500 proteins, 10% of which were also depleted in cells upon inhibitor treatment. Effect size and proportions were comparable to studies on USP7 inhibition (Steger *et al*, 2021). Whether the remaining 90% of ubiquitination sites are non-degradative events or did not result in protein degradation in the time-window studied, remains unclear. When we focussed on proteome changes after USP9X inhibition, we realised that USP9X seems to regulate distinct sets of proteins in a highly cell-type specific manner. This observation can at least partially explain the differences and lack of consistency and/or reproducibility of previously published results with regard to cellular roles of USP9X.

Importantly, we identify a core set of 17 proteins uniformly regulated by USP9X inhibition, across highly diverse cell lines. Our data suggests that these proteins represent a consistent set of direct targets of USP9X, and will be useful as direct proximal biomarkers for future studies. Indeed, proximal biomarker discovery has been a bane of DUB inhibitor research, and our multi-cell line approach may be more broadly applicable for other DUBs for which specific inhibitors are available.

The 17 commonly regulated proteins, in context of all proteins changed across all cell lines, allowed GO-term meta-analysis (Zhou *et al*, 2019) to unveil critical cellular processes regulated by USP9X inhibition, with the most prominent being the regulation of mitotic processes in metaphase. Other pathways included TGFβ signalling, translation, and regulation of cytoskeletal architecture. Here our data was consistent with a subset of previous studies that had shown through USP9X genetic deletion, regulation of cell cycle progression (Engel *et al*, 2016; Skowyra *et al*, 2018; McGarry *et al*, 2016; Vong *et al*, 2005), SMAD signalling (Dupont *et al*, 2009; Stegeman *et al*, 2013), and ribosomal quality control (Clancy *et al*, 2021). It remains unclear how the specificity of USP9X regulation of these pathways is provided. Does USP9X regulate specific protein pools and act as a rheostat to adjust the amounts of proteins required to establish signalling events or mitotic progression? Or, is USP9X located at hotspots within the cell where it regulates centrosome biogenesis or TGFβ signalling *in situ*? Previous data has shown USP9X appears to co-locate with centrosomal proteins, and has been located to specific sites in cells (Li *et al*, 2017; Wang *et al*, 2017; Reijnders *et al*, 2016; Murray *et al*, 2004), which may explain how USP9X appears to regulate e.g. multiple proteins in centrosomal complexes. However, it is also possible that specific loss of a critical protein leads to collateral destabilisation of larger complexes and indirect protein depletion. A further insight from our proteomic analysis is the identification of numerous E3 ligases, including HECTD1, DTX3L and MKRN2 within the top regulated proteins across cell lines, and numerous additional E3 ligases in individual cell lines, suggesting that USP9X may sit atop a hierarchy of UPS processes, with USP9X inhibition unleashing a cascade of secondary events that may induce further proteome changes. This observation may also account for the unique proteomic changes observed in individual cell lines upon USP9X inhibition that may be dependent on secondary processes.

Importantly, however, we confirm the role of USP9X in cell cycle regulation through our proteomic analysis and cell imaging studies, that show that WEHI-092 leads to mitotic arrest in cancer cells, which intriguingly was not associated with cell death. This explains observed phenotypes in cancer cell line panels where WEHI-092 leads to mild growth arrest but not death, with few exceptions, that is exaggerated in longer-term studies assessing clonogenic potential.

It was striking to note that UO-31 cells, a papillary renal cell carcinoma cell line, was highly sensitive to USP9X inhibition. In this cell line, USP9X appears to regulate multiple essential pathways. It will be interesting to further study this apparent USP9X dependency and understand whether the USP9X dependency exists in other specific cancer settings. Establishing WEHI-092 as an agent that arrests mitosis, could be beneficial in exploiting cancer vulnerabilities. Mitotic spindle agents such as taxol are often front-line chemotherapy treatments and can be boosted by co-treatment with secondary mitosis inhibitors or DNA damaging agents. It would be interesting to test USP9X inhibitors, which had little or no toxicity on primary cells, in combination therapy, which may sensitise cancer cells to chemotherapy.

Overall, our work exemplifies how specific DUB inhibitors can help unravel DUB biology and increase understanding of DUB function in diseases such as cancer. Broad cell line screening with high-quality compounds is becoming feasible, can reveal biomarkers, and also overcome limitations arising from genetic manipulation. Our work, together with previous complementary studies (Steger *et al*, 2021; Doherty *et al*, 2022; Pinto-Fernández *et al*, 2019; Varca *et al*, 2021; Chan *et al*, 2023), paves the way to streamline DUB drug discovery and refine target rationales for DUB inhibitors for human disease.

## Material and Methods

### Medicinal chemistry

Details on medicinal chemistry including synthesis of compounds used in this study can be found in the **Supplementary Material**.

### Protein biochemistry

#### Reagents

Ubiquitin Rhodamine110 (Ub-Rh110MP, UbiQ Bio, UbiQ-126), isopropyl-ß-D- thiogalactoside (IPTG, GoldBio I2481C100), 2-mercaptoethanol (Sigma-Aldrich M3148), imidazole (Sigma-Aldrich 56749), lysozyme (Glentham Life Science GE8228), DNase I (Roche 11284932001), Bovine serum albumin (Sigma-Aldrich, A9647), Triton™ X-100 (Sigma-Aldrich T9284), tris(hydroxymethyl)aminomethane (Tris, Sigma Aldrich 9210- OP), 2-propanol (Sigma-Aldrich, 1.09634.2511), tris(2-carboxyethyl)phosphine hydrochloride (TCEP, GoldBio TCEP25), dimethyl sulfoxide (DMSO, Sigma-Aldrich, 472301), sodium phosphate dibasic (Sigma-Aldrich, 04276), sodium phosphate monobasic (Sigma-Aldrich, S8282), 4-(2-hydroxyethyl)piperazine-1-ethanesulfonic acid (HEPES, Sigma Aldrich, H4034), Tween® 20 (Sigma-Aldrich, P7949).

#### Molecular biology

An expression construct for the USP9X catalytic domain (aa 1551- 1970, NCBI Reference Sequence: Q93008-1), was codon optimised for bacterial expression, synthesised (Integrated DNA Technologies) and cloned into the pOPIN-S vector (Berrow *et al*, 2007) which was digested with KpnI and HindIII using In-Fusion™ HD cloning (Takara Clontech). For SPR, constructs were ordered with a GGS linker followed by an N-terminal AviTag™ (aa sequence: GLNDIFEAQKIEWHE) and were cloned as above. For the USP9X Δβ catalytic domain a deletion construct ΔE1924-K1943 and for USP9X/USP24 chimera, a construct with USP9X Y1757-A1766 replaced with the USP24 (NCBI Reference Sequence: NP_056121) aa motif FVRGEVLEGS were used. All constructs were confirmed by Sanger sequencing (AGRF).

#### Protein expression and purification

Protein expression vectors were transformed into *E. coli* BL21 cells. Cells were grown in 2x YT medium at 37°C (200 rpm) until OD600 = 0.6 was reached. Protein expression was induced by adding IPTG to a final concentration of 0.4 mM. Cultures were grown overnight (o/n) at 16°C, 200 rpm. Next, cultures were harvested by centrifugation at 5,000 x g for 10 min at 4°C. Cells were lysed by sonication in lysis buffer (25 mM Tris (pH 8.0), 500 mM NaCl, 5% (v/v) glycerol) at pH 8.0 supplemented with 1 mM TCEP, EDTA-free protease inhibitor cocktail tablets (Roche), lysozyme (2 mg/mL) and DNase I (100 µg/mL). Lysates were cleared by centrifugation at 20,000 rpm for 30 min at 4°C. The cleared supernatant was filtered (0.45 µm) and was then incubated with Ni-NTA His resin (EMD Millipore). The resin was eluted with elution buffer (25 mM Tris (pH 8.0), 250 mM NaCl, 250 mM imidazole, 1 mM TCEP) at pH 8.0. The eluate was incubated with 0.1 mg/mL SENP1 protease (purified as described previously (Pruneda *et al*, 2016)) to cleave the His / SUMO tag in dialysis buffer (25 mM Tris (pH 8.0), 200 mM NaCl, 5 mM 2-mercaptoethanol) at pH 8.0 o/n at 4°C. The following day, the concentrated and tag-cleaved protein was incubated with Ni-NTA His resin and the eluate was collected for purification by ion-exchange chromatography using a Resource Q 6 mL column (Cytiva). Eluates were pooled and concentrated using a 10 kDa membrane filter (Merck). Protein was further concentrated and purified by size-exclusion chromatography (SEC) using a HiLoad 16/600 SuperDex 75 pg column (Cytiva) in SEC buffer (25 mM Tris (pH 8.0), 150 mM NaCl, 1 mM TCEP) at pH 8.0. The pure protein fractions were pooled, concentrated to a concentration of 12 mg/mL and flash-frozen in liquid nitrogen. Protein was stored at -80°C. Ubiquitin suicide probes were prepared as described previously (Ekkebus *et al*, 2013; Gersch *et al*, 2017).

#### Ub-Rho cleavage

USP9X catalytic domain (final concentration 7.8 nM) was pre-incubated with USP9X inhibitors in a 12-point, 1 in 3 titration for 30 min at ambient temperature and subsequently mixed with 100 nM Ub-Rh110MP substrate in assay buffer (20 mM Tris (pH 8.0), 0.01% (v/v) Triton X 100, 5 mM DTT, 0.03% (w/v) BSA) at pH 8.0 for 35 min at ambient temperature. The reaction was stopped through addition of 10 mM citric acid. The assay was performed in 384-well format with a total reaction volume of 16 µL. Data was collected using the ClarioSTAR plate reader (BMG Labtech) at an emission of 535 nm. Blank corrected values were normalised to DMSO control (100% activity remaining) and were used for IC50 calculation using GraphPad Prism v10.3 using the [inhibitor] vs. response analysis (four parameter fit) model. IC50 values presented in our study are the mean of three independent biological repeats unless indicated otherwise. **SPR:** Experiments were performed on a Biacore 8K+ instrument (Cytiva). USP9X catalytic domain with an N-terminal AviTag™ was immobilised in 10 mM HEPES, 150 mM NaCl at pH 7.4 on a Series S Sensor Chip SA (Cytiva) by coupling. Compounds were diluted from 10 mM stocks in DMSO into PBS-P+ running buffer (20 mM phosphate buffer (sodium phosphate dibasic 39% (v/v) and sodium phosphate monobasic 61% (v/v)), 150 mM NaCl, 0.05% (v/v) Tween 20 and 1 mM TCEP at pH 6.2). Running buffer was supplemented with 2% (v/v) DMSO. Multi-cycle kinetics were performed with 120 s associations and 600 s dissociations at 30 µL/min with no further regeneration. Binding constants were determined in Biacore insight evaluation software (v5.0.18) at steady state taking the median response over a 5 s window beginning at 10 s before the end of the association phase. Final KD values are the mean of three independent experiments. For inhibitor activity assays, USP9X catalytic domain (di-ubiquitin cleavage assay: 750 nM, ubiquitin-probe assays: 2.5 µM) was pre-incubated with USP9X inhibitors for 30 min at ambient temperature and subsequently mixed with di-ubiquitin (final concentration 2 µM) in assay buffer (25 mM Tris (pH 8.0), 150 mM NaCl, 10 mM DTT) at pH 8.0 or was mixed with ubiquitin-propargylamine (Ub-PA) / ubiquitin-vinyl sulfonate (Ub-VS) probes at a final concentration of 10 µM in assay buffer. For both assays, the mixture was incubated over a time course and the reaction was stopped by mixing samples with 2X LDS sample buffer (Invitrogen) supplemented with 3% (v/v) 2-mercaptoethanol. All protein samples were run on a 4-12% Bis-Tris SDS-PAGE gradient gel (Invitrogen) for 60 min at 140 V. The gel was stained with InstantBlue® (Abcam) and imaged.

#### WEHI-092 specificity DUB panel

WEHI-092 specificity was assessed using the commercial DUB*profiler*™ (DUBprofiler-FLEX SPT, v8) platform. WEHI-092 was supplied, and testing was performed at Ubiquigent Ltd (Dundee, UK) at 50 µM WEHI-092 in 2 technical replicates.

#### Mass spectrometry-based 3X-FLAG-Ub-VS DUB competition assay with cell extracts (DUB IP-MS)

MCF-7 cells have been profiled for active DUBs previously (Pinto-Fernández *et al*, 2019). Lysates were generated as described previously (Turnbull *et al*, 2017) using 30 s freeze-thaw cycles (thrice) in lysis buffer (50 mM Tris (pH 7.4), 5 mM MgCl2 x 6H2O, 0.5 mM EDTA, pH 8.0, 250 mM sucrose and 1 mM DTT) at pH 7.5. For experiments with crude cell extracts, 50 µg MCF-7 lysate was incubated with 50 µM WEHI-092 for 60 min at ambient temperature. A pre-treatment of cell lysates with 0.5 µg Ub-VS was included for a baseline DUB response to the Ub-VS probe. 0.1 µg 3X-FLAG-Ub-VS probe was added to samples and incubated for 5 min at 37°C. Incubation with the probe was optimised to minimise replacement of non-covalent inhibitor WEHI-092 by the covalent probe. 3X-FLAG-Ub-VS labelled DUBs were captured by incubation with anti- FLAG M2 affinity gel resin (Sigma, A2220) for 3.5 h at 4°C on a rotator. Samples were washed thrice with lysis buffer and eluted from the resin with lysis buffer supplemented with 1% (w/v) SDS following a 10 min incubation. Two consecutive elutions were pooled and samples were stored at -80°C until processing.

#### HDX-MS

Deuterium labelling of USP9X catalytic domain was performed as described previously (Asadollahi *et al*, 2023) at 20°C for periods of 0, 6, 60, 6000 s using a PAL Dual Head HDX Automation manager (Trajan/LEAP) controlled by the ChronosHDX software. 3 µL of USP9X catalytic domain (∼25 µM) was transferred to 55 µL of non- deuterated (50 mM potassium phosphate buffer at pH 7.4 in H2O) or deuterated (50 mM potassium phosphate buffer pD 7 in D2O) buffer and incubated for the respective time with 80 µM WEHI-092 or 30 µM FT709. Quenching was performed by adding 50 µL of the protein mix to 50 µL of quench buffer (50 mM potassium phosphate buffer (pH 2.3), 4 M guanidine hydrochloride and 0.1% (v/v) n-dodecylphosphocholine) at 1°C. For online pepsin digestion, 80 µL of the quenched sample was passed over an immobilised pepsin column (2.1 × 30 mm Enzymate BEH, Waters) equilibrated in 0.1% (v/v) formic acid (FA) in H2O at 100 µL/min. To further reduce peptide carryover, n-octyl-β-d-glucopyranoside at 1% (w/w) was added to the pepsin column wash solution (1.5 M guanidine hydrochloride, 4% (v/v) acetonitrile (ACN), 0.8% (v/v) FA). Proteolysed peptides were captured and desalted by a C18 trap column (VanGuard BEH; 1.7 μm; 2.1 × 5 mm; (Waters)) and eluted with ACN and 0.1% (v/v) FA gradient (5% to 40% in 8 min, 40% to 95% in 0.5 min, 95% 1.5 min) at a flow rate of 80 μL/min and separated on an ACQUITY UPLC BEH C18 analytical column (1.7 μm, 1 × 50 mm, (Waters) delivered by ACQUITY UPLC I-Class Binary Solvent Manager (Waters)). For MS, an ion mobility equipped SYNAPT G2-Si mass spectrometer (Waters) was used. Instrument settings were: 3.0 KV capillary and 40 V sampling cone with source and desolvation temperature of 100°C and 40°C respectively. The desolvation and cone gas flow was at 80 L/h and 100 L/h, respectively. High energy ramp trap collision energy was from 20 to 40 V. All mass spectra were acquired using a 0.4 s scan time with continuous lock mass (Leu-Enk, 556.2771m/z) for mass accuracy correction. Data were acquired in HDMS^E^ (ion mobility) mode and peptides from non-deuterated samples were identified using Protein Lynx Global Server (PLGS, v3.0, Waters). To ensure high peptide selection stringency, we applied additional filter constraints of 0.3 fragments per residue, minimum consecutive product of 1, minimum score of 6, minimum intensity of 2500, maximum MH+ error of 5 ppm, retention time RSD of 10% and file threshold of 3 out of 6 HDMS^E^ files. The deuterium uptake values were calculated for each peptide using DynamX 3.0 (Waters). Deuterium exchange experiments were performed in technical triplicate for each of the timepoints. Peptides with a statistically significance difference in HDX were determined using the Deuteros (v2.0) (Lau *et al*, 2021) software with a hybrid significance test with a 99% confidence interval.

#### Multiple sequence alignment USP catalytic domains

Annotated catalytic domain sequences (USP9X UniProt ID Q93008-1, aa 1557-1956, USP24 UniProt ID Q9UPU5, aa 1689-2042, USP34 UniProt ID Q70CQ2, aa 1894-2239, USP28 UniProt ID Q96RU2, aa 162-650, USP25 UniProt ID Q9UHP3, aa 169-657) were extracted and a multiple sequence alignment (MSA) was performed using Clustal Omega (EMBL-EBI (Madeira *et al*, 2024).

#### AlphaFold3

For the AlphaFold3 (Abramson *et al*, 2024) models of USP9X and USP24 the following sequences were used: Ubiquitin (UniProt ID P0CG48, aa 1-76), USP9X catalytic domain with ubiquitin (UniProt ID Q93008-1, aa 1549-1970), USP24 catalytic domain with ubiquitin (UniProt ID Q9UPU5, aa 1688-2042), USP24 full-length (UniProt ID Q9UPU5, aa 1-2620). Source code was downloaded and run on internal servers.

### Studies in cells

#### Reagents

DMEM (Gibco, 11885-084), RPMI (in-house), Trypsin (Sigma), GlutaMAX™ (Gibco, 35050061), FBS (Bovogen, SFBS-F, S00JF), Horse serum (Thermo Fisher Scientific, 16050122), Insulin (Merck, I5500), Human EGF (Sigma, E9644), Hydrocortisone (Sigma, H0396), Propidium iodide (Sigma, P4170). Reagents include antibodies used for Western Blotting against USP9X (Bethyl, A301-350A), CEP55 (Cell Signaling Technology, 81693), MKRN2 (Abcam, ab72055), β-Actin (Santa Cruz Biotechnology, sc-47778 HRP), EDF1 (Abcam, ab174651), and secondary antibody against rabbit IgG coupled to horseradish peroxidase (Jackson ImmunoResearch, 111- 035-003).

#### Cell lines

MDA-MB-231, MiaPaca2, UO-31, HMEC-1, SK-MEL-2, OVACR8, DLD-1 and A549 cells were validated at CellBank Australia. Primary cells HUVEC (STEMCELL Technologies) and HDF (Lonza) were sourced from commercial providers. All other cell lines were sourced in-house. Cell lines were screened monthly for *mycoplasma* contamination using the MycoAlert® kit (Lonza LT07-318) as per the manufacturer’s instructions. All cells used were *mycoplasma* free. Details on culturing media for cell lines used in this study can be found in the **Supplementary Material**. All cell lines were cultured at 37°C, 5% CO2 in a standard tissue culture incubator. Human dermal fibroblasts were cultured at 37°C, 10% CO2.

#### Colony formation assays

Cells were seeded at low density (200 – 4,000 cells per well) after initial optimisation in a 6-well tissue culture treated plate (Falcon). 24 h post seeding, media was replaced, and compound was added. Cells were left to grow into colonies between 7-14 d depending on the cell line. For analysis, media was removed, and cells were fixed with 10% (w/v) formalin for 10 min. Colonies were stained with crystal violet (Sigma, 0.5% (w/v) in 20% (v/v) MeOH) for 10 min and washed thrice with distilled H2O. Plates were imaged using the ChemiDoc imaging system (BioRad) and analysed using the ImageJ ColonyArea plugin as described previously (Guzmán *et al*, 2014). **Live cell imaging assays:** Cells were seeded at varying density (1,000 – 8,000 cells per well) after initial optimisation in a 96-well tissue culture treated plate (Thermo). Cells were incubated o/n, and the next day, inhibitors and controls diluted in cell media were added to the cells to a final volume of 200 μL per well. Cell death was measured using propidium iodide (PI, Sigma). Cells were imaged with 4 images per well every 2 h for up to 96 h using the IncuCyte live-cell analysis system (S3, Sartorius). Cell proliferation data was analysed using Microsoft Excel by normalising confluence to control conditions. For cell killing readouts, the number of PI positive cells was normalised to the control conditions for each timepoint.

#### NCI cancer cell line panel

NCI-60 cancer cell panel (National Cancer Institute, US) was used to assess cell line sensitivity to WEHI-092 and WEHI-680 treatment. WEHI-092 and WEHI-680 were supplied and experiments were performed at the NCI developmental program (DTP). WEHI-092 was tested at 10 µM. Data shown is the mean of two technical replicates and is shown relative to the no-drug control, and relative to the time zero number of cells.

#### Immunoblotting

Cells were plated in 24-well tissue culture treated plates (Falcon), and one day post plating media was replaced, and cells were treated with WEHI-092. The next day, cells were lysed in DISC lysis buffer (20 mM TRIS/HCl (pH 7.5), 150 mM NaCl, 2 mM EDTA (pH 8.0), 1% (v/v) Triton X 100, 10% (v/v) glycerol) supplemented with 2% (w/v) SDS and PhosSTOP tablets (Roche). To shred DNA, DISC lysates were run through polypropylene columns (Pierce). Proteins were separated by SDS-PAGE on a 4-12% Bis- Tris gradient gel (Invitrogen) for 10 min at 70 V followed by 90 min at 120 V. Samples were transferred to Immobilon-E Transfer PVDF membranes (Merck) by wet transfer at 4°C for 60 min at 100 V. Membranes were blocked in 5% (w/v) skim milk (Devondale) in TBS-T (TBS supplemented with 0.1% (v/v) Tween-20) for 60 min at ambient temperature. Next, membranes were washed twice in TBS-T and incubated with primary antibodies at 4°C o/n. Primary antibodies were diluted in antibody dilution buffer (5% (w/v) BSA (pH 5.2, Sigma) in TBS-T with 0.04% (v/v) sodium azide) as per the manufacturer’s instructions. Membranes were washed thrice for 5 min each in TBS-T. Next, secondary anti-rbIgG antibody (1 in 10,000, Jackson ImmunoResearch; Cat# 111-035-003) conjugated with HRP diluted in 5% (w/v) skim milk in TBS-T was incubated with the membranes for 60 min at ambient temperature. β-actin antibody conjugated to HRP (Santa Cruz; Cat# sc-47778 HRP) was used at a 1 in 20,000 dilution in 5% (w/v) skim milk in TBS-T for 40 min at ambient temperature as a loading control. Next, membranes were washed four times for 5 min each in TBS-T. Membranes were developed in ECL (BioRad) and imaged using the ChemiDoc imaging system (BioRad). Images were processed using Image Lab software.

#### Protein extraction and digestion (ubiquitinomics)

Cells were harvested directly off the cell culture plates using pre-heated sodium deoxycholate (SDC) buffer (1% (v/v) SDC, 10 mM TCEP, 40 mM 2-chloracetamide (CAA), 75 mM Tris-HCl at pH 8.5). Samples were incubated at 85°C for 10 min and allowed to cool to ambient temperature prior to the addition of universal nuclease (Thermo) and incubated for 5 min prior to centrifugation and the clarified lysates were collected. Protein concentrations were determined using the Pierce™ BCA Protein Assay Kit (Thermo) and proteins were digested with Lys-C (Wako, 129–02541) and TPCK-treated Trypsin (Thermo Cat# 20233) mix o/n at 37°C with a 1:50 enzyme to protein ratio. As described previously, the digestion was stopped by adding three volumes 1% (v/v) trifluoroacetic acid (TFA, Sigma) in 2-propanol and loaded onto SDB-RPS cartridges (Strata™-X-C, 100 mg, Phenomenex Inc.) (Hansen *et al*, 2021). Columns were activated and pre-equilibrated with 3 mL of 30% (v/v) MeOH / 1% (v/v) TFA and washed with 3 mL of 0.2% (v/v) TFA. Samples were loaded and washed twice with 3 mL 1% (v/v) TFA in 2-propanol and once with 3 mL 0.2% (v/v) TFA / 2% (v/v) ACN. Peptides were eluted twice with 2 mL of 1.25% (v/v) NH4OH / 80% (v/v) ACN and diluted with H2O to a final ACN concentration of 30% (v/v). The eluates were snap-frozen in liquid nitrogen and lyophilised o/n.

#### Crosslinking of K-GG antibody

Crosslinking was performed as described previously (Udeshi *et al*, 2013). Briefly, 2.5 μL IgG Sepharose 4B (Cytiva) was incubated with 31 µg of an in-house K-GG antibody for 1 h at ambient temperature. Antibody-bound beads were washed thrice with 100 mM sodium tetraborate (pH 9.0) and then crosslinked for 30 min at ambient temperature with 0.5 mL of 20 mM dimethyl pimelimidate in 100 mM sodium borate (pH 9.0). The cross-linking reaction was then quenched by the addition of 200 mM ethanolamine (pH 8.0) and washed twice before being incubated for 2 h with 200 mM ethanolamine (pH 8.0). The beads were washed thrice with IAP buffer (50 mM MOPS pH 7.2, 10 mM Na2HPO4, 50 mM NaCl).

#### K-GG peptide enrichment and LC-MS/MS sample preparation

K-GG peptide enrichment was performed as described with some minor modification (Udeshi *et al*, 2013). Briefly, peptides were resuspended in 0.5 mL of cold IAP buffer and incubated with 2.5 µL of packed cross-linked K-GG antibody-bead conjugate (corresponding to 31 µg of antibody per sample) for 2 h at 4°C with end-over-end rotation. Beads were transferred to glass-fibre filter stage tips and washed twice with 1 mL of IAP buffer and an additional three times with cold H2O via centrifugation between each wash (Hansen *et al*, 2021). The glass-fibre filter stage tips were then stacked onto SDB-RPS stage tips. The SDB- RPS stage tips were pre-activated and equilibrated with the addition of 60 µL of 2- propanol, 60 µL 80% (v/v) ACN and 100 µL 0.2% (v/v) TFA with centrifugation occurring between each addition. K-GG peptides were eluted off the beads and directly captured onto the SDB-RPS stage tips via two separate 100 µL of 0.15% (v/v) TFA elutions and centrifugation. Peptides were then desalted by washing the SDB-RPS stage tips with 0.2% (v/v) TFA / 2% (v/v) ACN, prior to elution with 60 µL 80% (v/v) ACN / 2.5% (v/v) NH4OH directly into level 3 SureStart 0.2 mL MS vials (Thermo). Peptides were Speedvac dried and then resupended in 10 µL of 0.1% (v/v) FA / 2% (v/v) ACN, with 4 µL injected into the mass spectrometer.

#### Protein extraction and digestion

For global proteomics, cell pellets from one well of a 6-well plate per technical replicate (n=5) were lysed in 200 µL of preheated (95°C) buffer (2.5% (v/v) SDS in 100 mM Tris-HCl, pH 8.5). DNA was hydrolysed with the addition of 2 µL neat TFA and lysates were neutralised to pH 8.5 by addition of 1 M Tris-HCl as previously described (Dagley *et al*, 2019). Protein concentration was determined using Pierce™ BCA Protein Assay Kit (Thermo) following manufacturer’s instructions. For DUB IP-MS, samples were processed using S-Trap spin columns (Profiti) by following the manufacturer’s instructions. Briefly, samples were reduced with 5 mM TCEP and alkylated by addition of 20 mM CAA with incubation at 55°C for 15 min each prior to addition of 2.5% (v/v) phosphoric acid. In-column digestion was performed with a Lys-C (Wako, 129– 02541) and SOLu-Trypsin (Sigma-Aldrich, EMS0004) at 1 µg per column for 1:45 h at 47°C. For global proteomics, cell lysates (20 µg protein per replicate) were transferred to 0.5 mL LoBind deep well plate (Eppendorf) prepared for MS analysis using the modified SP3 protocol (Hughes *et al*, 2019), with some modifications. Briefly, samples were subjected to simultaneous reduction and alkylation with a final concentration of 10 mM TCEP and 40 mM CAA followed by heating at 95°C for 10 min. Prewashed magnetic PureCube Carboxy agarose beads (20 µL, Cube Biotech) were added to all the samples along with ACN (70% (v/v) final concentration) and incubated at ambient temperature for 20 min. Samples were placed on a magnetic rack and supernatants were discarded, and beads were washed twice with 70% (v/v) ethanol and once with neat ACN. ACN was completely evaporated from the tubes using a CentriVap (Labconco) before the addition of digestion buffer (50 mM Tris-HCl, pH 8) containing 1 µg each of enzymes Lys-C (Wako, 129–02541) and SOLu-Trypsin (Sigma-Aldrich, EMS0004). Trypsin-LysC on-bead digestion was performed with agitation (400 rpm) for 1 h at 37°C on a ThermoMixer C (Eppendorf). For DUB IP-MS and global proteomics, the samples were transferred to pre- equilibrated C18 StageTips (2× plugs of 3M Empore resin, no. 2215) following digestion for sample clean-up. The eluates were lyophilised to dryness before being reconstituted in 150 µL 0.1% (v/v) FA / 2% (v/v) ACN ready for MS analysis.

#### LC-MS/MS measurements

Peptides were loaded on a 15 cm C18 fused silica column with an integrated emitter tip (IonOpticks, ID 75 µm, OD 360 µm, 1.6 µm C18 beads) which was maintained at 50°C using a column oven. <u>DUB IP-MS</u> was performed on a Neo Vanquish (ThermoFisher) directly coupled online with the mass spectrometer (Eclipse ThermoFisher) and peptides were separated with a binary buffer system of buffer A (0.1% (v/v) FA) and buffer B (99.9% (v/v) ACN plus 0.1% (v/v) FA), at a flow rate of 250 nL/min. The gradient started at 2% B and increased to 34% B in 80 min before increasing to 85% B within 3 min and held for 15 min prior to returning to 2% B and re- equilibration. The mass spectrometer was operated in positive polarity mode with a capillary temperature of 275°C. The data-independent acquisition (DIA) methods consisted of a MS1 scan (*m*/*z* = 350-1,650) with an automatic gain control (AGC) target of 1.2 × 10^6^ and a maximum injection time of 60 ms (R = 120,000). DIA scans were acquired at R = 30,000, with an AGC target of 3 × 10^5^, “auto” for injection time and a default charge state of 3. The spectra were recorded in profile mode and the stepped collision energy was 25, 27.5, 30% normalised collision energy. 44 non-uniform DIA segments were set to achieve an average of 5 data points per peak. <u>K-GG samples</u> were acquired with a Neo Vanquish (Thermo) directly coupled online with an Astral mass spectrometer (Thermo) and peptides were separated with a binary buffer system of buffer A (0.1% (v/v) FA) and buffer B (80% (v/v) ACN plus 0.1% (v/v) FA), at a flow rate of 400 nL/min. The gradient started at 2% B and increased to 34% B in 30 min before increasing to 100% B within 0.1 min and held for 3 min prior to returning to 2% B and re- equilibration. The mass spectrometer was operated in positive polarity mode with a capillary temperature of 275°C. The DIA methods consisted of a MS1 scan (*m*/*z* = 380- 980) with an AGC target of 5 × 10^6^ and a maximum injection time of 5 ms (R = 240,000). DIA scans were acquired with the Astral detector with an AGC target of 8 × 10^4^ and 3 ms maximum time. Fragmentation occurred in the higher-energy collisional dissociation (HCD) cell with a normalised stepped collision energy was 25% and the spectra were recorded in profile mode. 199 non-uniform DIA windows across 380-980 were collected with a maximum injection time of 3 ms and a 0.6 s loop control which achieved an average of 5 data points per peak. <u>Global proteomic samples</u> were acquired using a custom nano-flow high-performance liquid chromatography (HPLC) system (Thermo Ultimate 300 RSLC Nano-LC, PAL systems CTC autosampler). The HPLC was coupled to a timsTOF Pro (Bruker) equipped with a CaptiveSpray source. Peptides were loaded directly onto the column at a flow rate of 600 nL/min with buffer A (99.9% Milli-Q H2O, 0.1% (v/v) FA) and eluted at 400 nL/min on a 30 min linear analytical gradient of increasing buffer B (90% (v/v) ACN, 0.1% (v/v) FA) from 2 to 34%. The timsTOF Pro MS was operated in diaPASEF mode using Compass Hystar 5.1. The settings on the thermal ionization MS (TIMS) analyser were as follows: Lock Duty Cycle to 100% with equal accumulation and ramp times of 100 ms, and 1/K0 Start 0.6 Vs/cm^2^ End 1.6 Vs/cm^2^, Capillary Voltage 1400 V, Dry Gas 3 L/min, Dry Temp 180°C. The DIA methods were set up using the instrument firmware (timsTOF control 2.0.18.0) for data-independent isolation of multiple precursor windows within a single TIMS scan. The method included two windows in each diaPASEF scan, with window placement overlapping the diagonal scan line for doubly and triply charged peptides in the m/z – ion mobility plane across 16 × 25 m/z precursor isolation windows (resulting in 32 windows) defined from m/z 400 to 1,200, with 1 Da overlap, and collision induced dissociation (CID) collision energy ramped stepwise from 20 eV at 0.8 Vs/cm^2^ to 59 eV at 1.3 Vs/cm^2^.

#### Raw data searching and analysis

MS raw files were processed using DIA-NN 1.8.1 as described previously (Steger *et al*, 2021). Global and DUB IP-MS raw files were searched with the following settings: Trypsin specificity, peptide length of 7-30 residues, cysteine carbamidomethylation enabled as a fixed modification, variable modifications set to N- terminal protein acetylation and oxidation of methionine, the maximum number of missed cleavages at 1 (DUB IP-MS) and 2 (global proteomics) and match between runs (MBR) enabled within individual cell lines. For DUB IP-MS, modification UniMod:35 with mass delta 15.9949 at M were considered as variable.

For ubiquitinomics, library-free searching with K-GG variable modification enabled with a maximum of two modifications and two-missed cleavages was performed with the reviewed human proteome (Uniprot), with MBR enabled and Robust LC quantification.

Precursors were consolidated by retaining the precursor that was detected in the most number of samples and had the highest intensity. Data were filtered for K-GG peptides that were detected in four out of five samples in at least one condition (unless specified otherwise) and differential expression analysis performed using the DEP R package (Zhang *et al*, 2018). Data was normalised with variance stabilising normalisation (VSN) and missing values imputed using a mixed Bayesian principal component analysis (BPCA) and Min method (unless stated otherwise in the figure legends). Significance testing of log2-transformed intensities was performed, and Benjamini-Hochberg corrected *p*-values < 0.05 were considered as significant. K-GG fold changes were corrected for changes in the proteome unless stated otherwise.

Downstream processing and differential expression analysis was performed in R using limma (Ritchie *et al*, 2015) and DEP packages. For DUB IP-MS and global proteomics quantities were determined using MaxLFQ (Cox *et al*, 2014a) for proteins with at least two peptides, data were filtered for proteins that were observed in at least four out of five replicates in at least one condition. VSN normalised and imputed using missing values imputed using a mixed BPCA and MinD method. For DUB IP-MS, Benjamini-Hochberg corrected *p*-values < 0.05 were considered as significant and DUBs were considered responders if at least 20% capture was lost upon Ub-VS pre-treatment. For global proteomics samples, significance testing of log2-transformed intensities was performed, and Q-values from limma (also called p.adj in our figure legends) < 0.05 were considered significant.

#### Lattice light sheet imaging

MDA-MB-231, MiaPaca2, and HMEC-1 cells were seeded at 2-15 x 10^4^ cells / mL in 8-well glass µ-slides (ibidi). The next day, cells were treated with 15 µM WEHI-092 / DMSO. SPY650-Tubulin, SPY555-DNA (both Spirochrome, Inc.), and Annexin V FITC (Sigma) were added at a final concentration of 1 in 1,000. Cells were left to grow for 24 h before time-lapse live-cell data was acquired using a Lattice Light Sheet 7 (Zeiss). Light sheets (488 nm, 561 nm and 640 nm) of 30 µm in length with a thickness of 1 µm were created at the sample plane via a 13.3X 0.44 numerical aperture (NA) objective. Fluorescence emission was collected via a 44.83X 1 NA detection objective via a multiband stop, LBF 405/488/561/633, filter. Aberration correction was set to a value of 170 to minimize aberrations as determined by imaging the Point Spread Function using 100 nm fluorescent microspheres. Data were collected with a frame time of 10 ms and a z step of 0.4 μm with 834 frames per 290 μm by 290 μm. Individual regions were imaged in parallel across six wells of the eight well chamber slide. Data are presented as Maximum Intensity Projections (MIPs) and these MIPs were used for all downstream analysis. Tile regions were imaged for 24 h with 10 min intervals at 37°C, 5% CO2. Analysis was performed using the MIPs of the different full fields of cells over the 24-48 h time period. Mitosis and mitosis arrest was classified through a combination of automated image analysis and manual validation. MIPs were processed using the TrackMate plugin in ImageJ/Fiji (Schindelin *et al*, 2012; Ershov *et al*, 2022). Segmentation was performed using the pre-defined StarDist model (Schmidt *et al*, 2018) on the SPY555-DNA channel and followed by an auto-threshold using the “Quality” filter. Small erroneous detected spots were further removed through a manually adjusted size filter. The remaining segmented nuclei were tracked using an Advanced Kalman Tracker capable of splitting and merging tracks which is necessary for tracking dividing cells. An initial search radius of 30 µm followed by a further search radius of 20 µm and a max frame gap of 2 were set. The mean intensity of all tracks was then measured to determine mitosis and mitosis arrest. Mitosis arrest was defined when a cell’s nuclei increased in mean intensity but did not progress through to division and remained with a high mean intensity value, indicating condensed chromatin. Successful mitosis was defined by cells whose mean intensity increased leading into mitosis but was followed by a splitting in the subsequent tracks. All identified mitosis and mitosis arrest events were manually validated and the frame number indicating the entry of mitosis was noted for all events. All videos were generated using ImageJ/Fiji.

## Supporting information

Combined Supplementary Data

## Acknowledgments

The authors would like to thank present and past members of the Ubiquitin Signalling Division at WEHI. We thank Sylvie Urbé and Michael J. Clague (University of Liverpool) for critical comments on the manuscript, and Kum Kum Khanna (Mater Research, Queensland) for early discussion on CEP55. We thank the National Drug Discovery Centre (WEHI, Parkville) for early compound testing. HDX-MS data was collected at the Bio21 proteomics facility. Proteomics data were collected at the WEHI proteomics facility. Imaging data was collected at the WEHI Centre for Dynamic Imaging. NCI-60 cancer cell panel testing was performed at the NCI-DTP. Schematics were created in BioRender under the license: Schenk, P. (2025) https://BioRender.com/kjpfiep.

The National Drug Discovery Centre is supported by the Australian Government Medical Research Future Fund (MRFF) Grant ID EPCD000033, and the Victorian Government. The laboratory of RF is supported by The Galbraith Family Charitable Trust, the K & M Foundation for Women, the Betty Deller King Bequest, the Rae Foundation and the Berwick opportunity shop. This work has been supported by an NHMRC Investigator Grant GNT117812 to DK.

## Author contributions

DK conceived the project. DK, APN, and RF jointly supervised the project and conceptualised experiments with contribution from PJAE. RF and DK secured funding for this project. PS performed all experiments and analysed all data unless indicated otherwise. SMD undertook medicinal chemistry studies. PS, DHM, VV and SAC performed ubiquitinomics and global proteomics experiments with contribution from LFD. PS and C-SA performed HDX-MS with contribution from NW. PS and NG performed lattice light sheet imaging and NG analysed the data. DJC contributed to DUB-IP-MS. BGCL, TAK and KNL contributed to initial compound screening. PS and DK wrote the manuscript, which was edited by PJAE and other authors.

## Competing interests

DK is founder, shareholder and SAB member of Entact Bio and Proxima Bio, and co-founder and SAB member of Ternarx. RF is co-founder and scientific lead at Ternarx.

## Materials and Correspondence

Requests for materials and correspondence should be addressed to DK.

## Data Availability

All data is available within this paper or as **Supplementary Material**. **Source Data** is available for all Figures of the manuscript.

The MS proteomics data have been deposited to the ProteomeXchange Consortium via the PRIDE (Perez-Riverol *et al*, 2024) partner repository.

## Notes

### Competing Interest Statement

The authors have declared no competing interest.

