## Supplementary material for "Global analysis of cancer cell responses to USP9X inhibition": Combined Supplementary Data

Philipp Schenk *et al*.

This file includes:

Fig. S1 to S8

Movies S1 to S4 legends

Table S1

Medicinal Chemistry

Other Supplementary Material for this manuscript include the following:

Movies S1 to S4

Figure S1

A

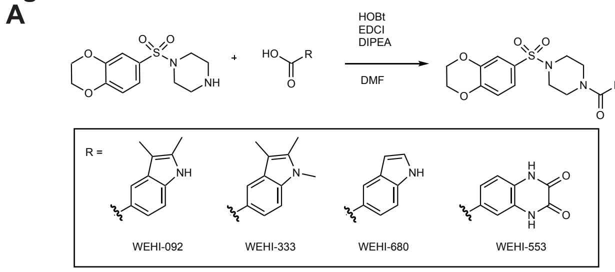


B

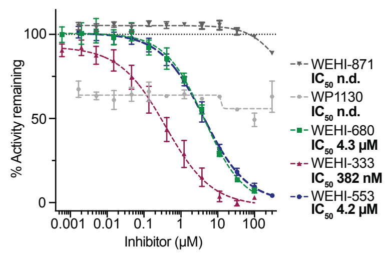


D

| Compound | Steady State Affinity ($K_D$) [µM] | $IC_{50}$ [µM] |
|---|---|---|
| FT709 | 0.069 ± 0.002 (N=3) | 0.124 (N=4) |
| **WEHI-092** | **1.05 ± 0.20 (N=3)** | **0.254 (N=4)** |
| WEHI-333 | 1.05 ± 0.26 (N=3) | 0.382 (N=3) |
| WEHI-680 | 3.51 ± 0.58 (N=3) | 4.3 (N=3) |
| WEHI-871 | n.d. (N=3) | n.d. (N=3) |
| WEHI-553 | Not tested | 4.2 (N=6) |

C

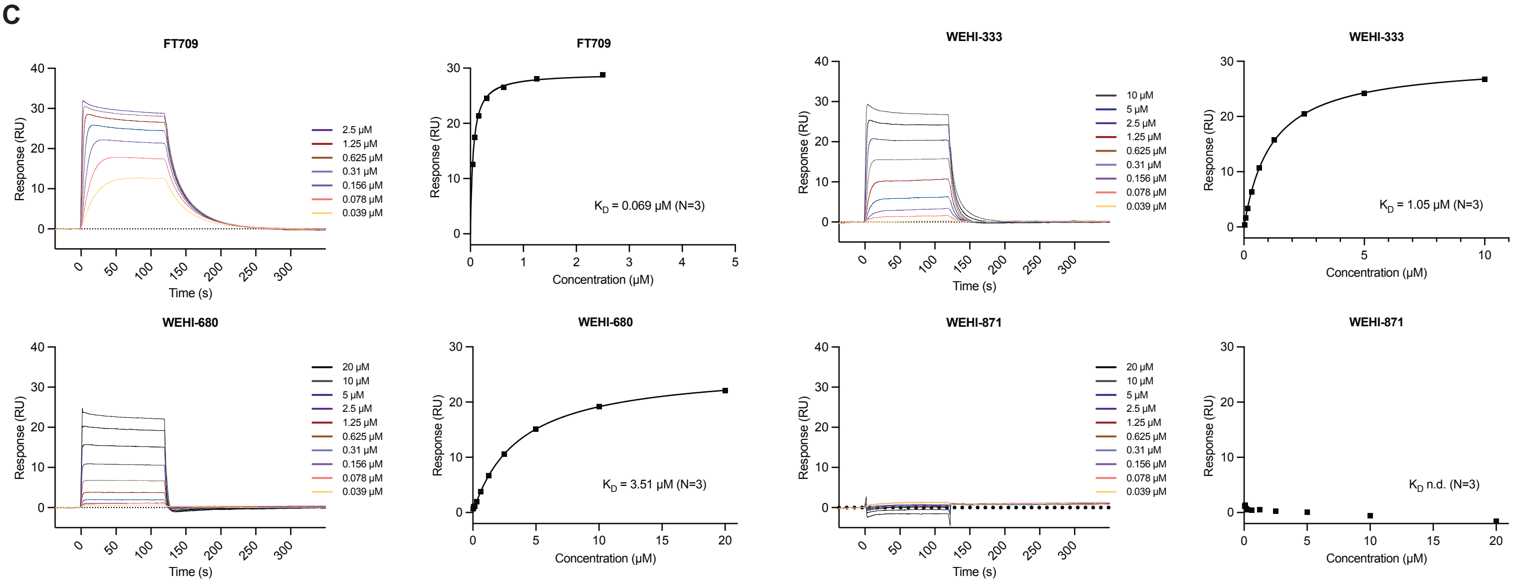


E

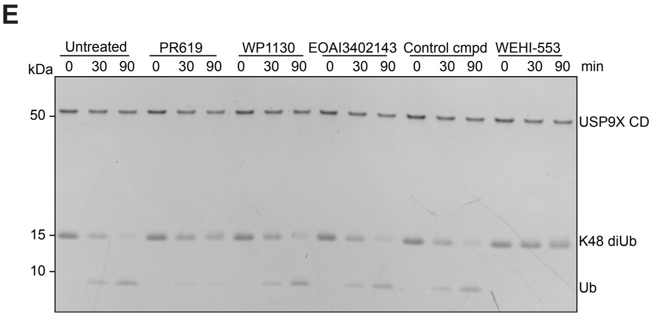


F

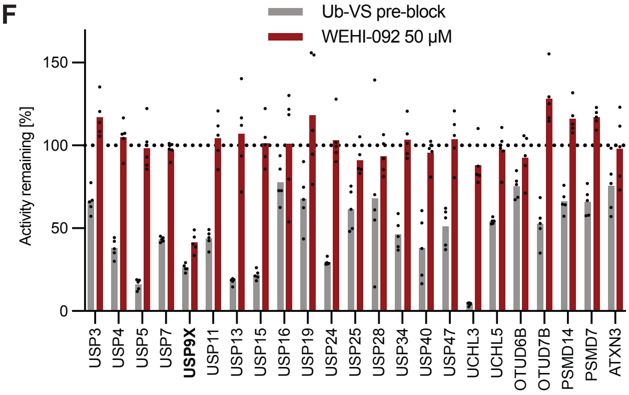

**Fig. S1. WEHI series USP9X inhibitors chemistry and assessment.**

(**A**) Chemical synthesis of WEHI series USP9X compounds. See **Supplementary Material: Medicinal Chemistry** for further details. (**B**) Ub-Rho cleavage assay to assess inhibitor potency against recombinant USP9X catalytic domain. Data shown is the mean of three independent biological repeats with two technical replicates per experiment. WP1130 is the mean of two independent biological repeats. A nonlinear curve fit generated using GraphPad Prism (v10.3) was used for calculating $IC_{50}$ values. n.d. – not determined; error bars, SEM. (**C**) SPR binding assay for USP9X inhibitors to bind the USP9X catalytic domain. For each compound: *Left*, fitted binding curve; *Right*, raw sensorgram data. $K_D$ shown is the mean of three biological repeats. Representative graphs from one experiment are shown. (**D**) Overview table of $K_D$ (determined via SPR) and $IC_{50}$ (determined via Ub-Rho assay) for compounds tested in this study (n.d. – not determined). (**E**) Coomassie gel of a K48-linked diUb cleavage assay with 150 µM compound. Representative gel of two independent biological repeats. (**F**) DUB IP-MS experiment to assess maximum level of inhibition with Ub-VS probe. Pre-treatment of MCF-7 cell lysates with 0.5 µg Ub-VS for 20 min at ambient temperature. Data shown is the mean of five biological replicates which are shown as individual datapoints.

### Figure S2

A

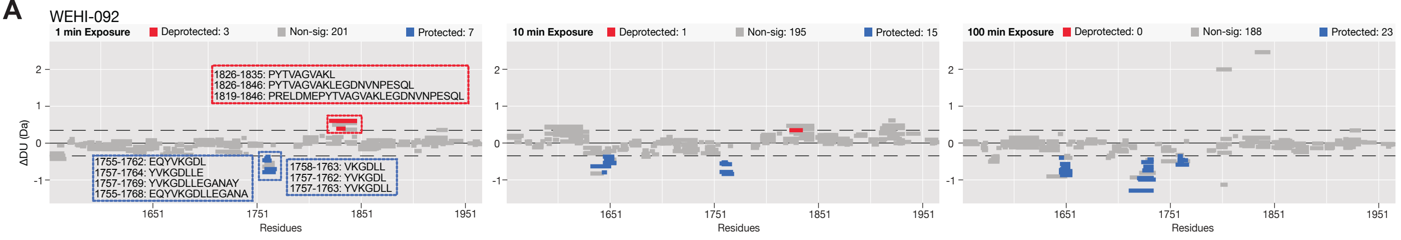


B

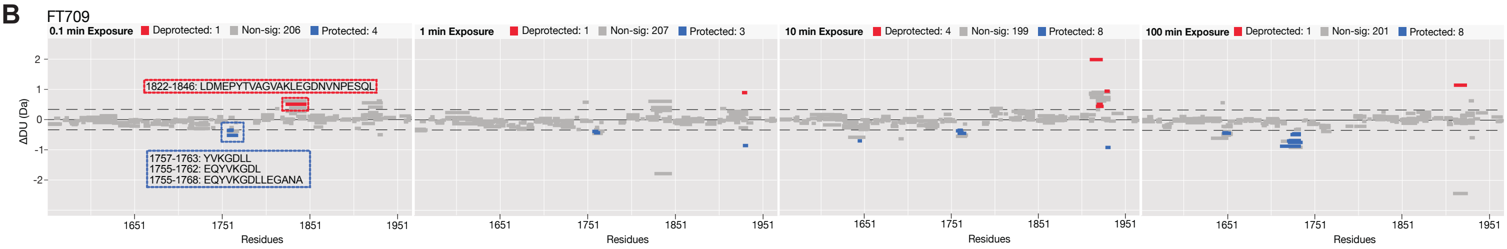


C

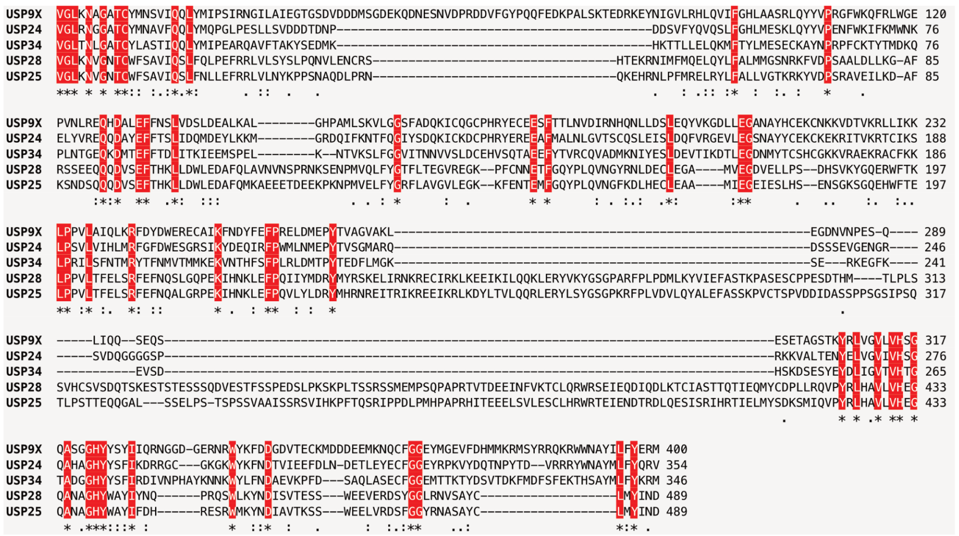


D

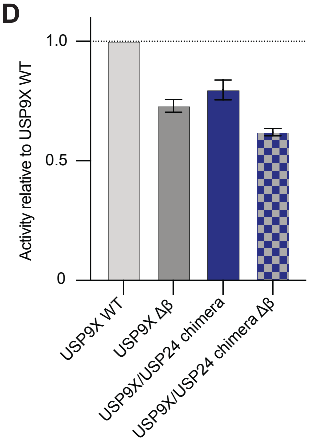


E

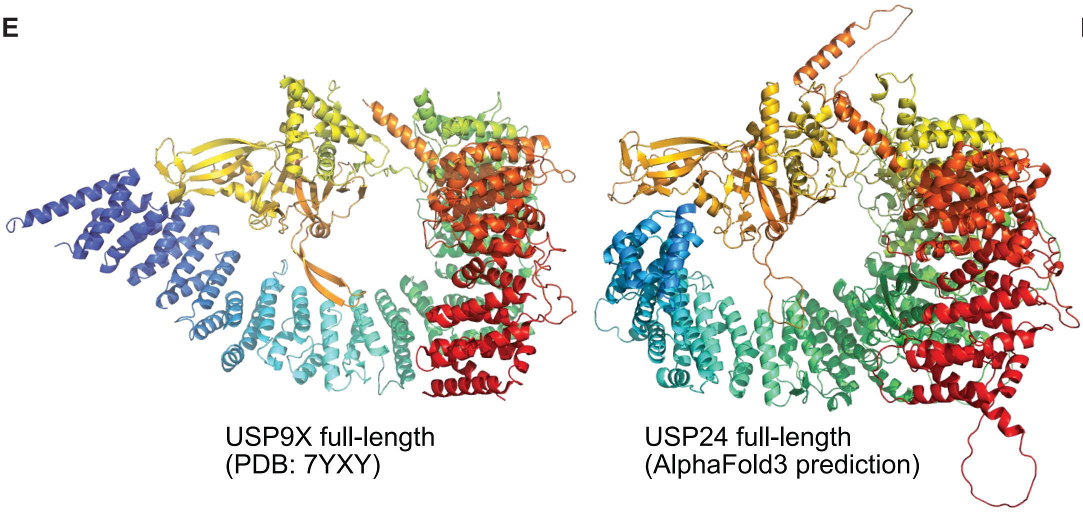


F

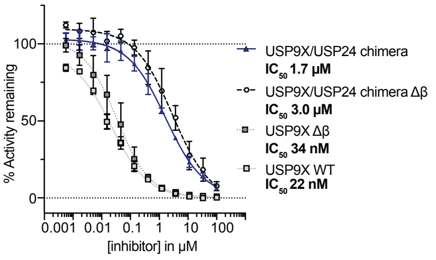

**Fig. S2. HDX-MS and USP9X catalytic domain mutants.**

(**A**) Hybrid Woods differential plot highlighting significant ($p < 0.01$) peptide differences in hydrogen-deuterium uptake between USP9X catalytic domain apo and WEHI-092 bound protein protected peptides in blue and deprotected peptides in red. The dashed lines indicate the confidence limit at 99% (hybrid significance test). Plots were generated using Deuteros software (v2.0). (**B**) As in **A** and **Fig. 2A**, but for FT709. (**C**) Sequence alignment of the indicated human USP DUB catalytic domains. (**D**) Catalytic activities of indicated wild-type (WT) and mutant USP9X proteins, assessed by Ub-Rho cleavage assays. Data shown is the mean of three independent biological repeats with four technical replicates per experiment. Error bars, SEM. (**E**) Side by side comparison of full-length USP9X (PDB: 7YXY) and full-length USP24 (AlphaFold3 prediction, residues 1-400 removed for visualisation purposes). (**F**) FT709 potency against recombinant USP9X catalytic domain variants: USP9X wild-type (WT), USP9X/USP24 chimera (USP9X with five USP24-like mutations in compound binding region), USP9X $\Delta\beta$ (USP9X $\beta$-hairpin deletion) and USP9X/USP24 chimera $\Delta\beta$ (USP9X $\Delta\beta$ with five USP24-like mutations) determined from Ub-Rho cleavage assay. Data shown is the mean of two independent biological repeats with two technical replicates per experiment. Curve shown is the nonlinear curve fit generated using GraphPad Prism (v10.3) and was used for calculating the $IC_{50}$ values. Error bars, SEM.

### Figure S3

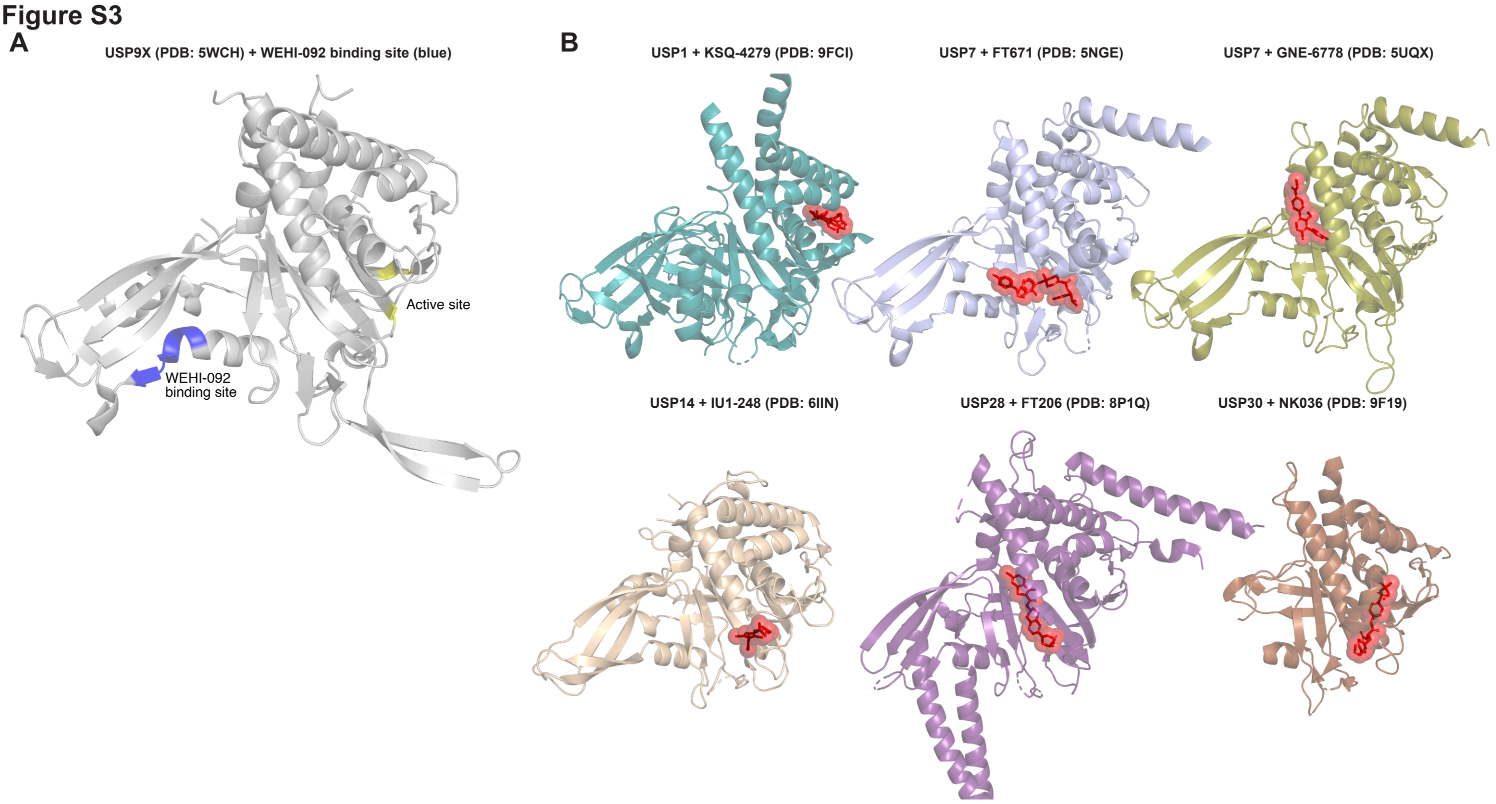
A
USP9X (PDB: 5WCH) + WEHI-092 binding site (blue)
Active site
WEHI-092
binding site
B
USP1 + KSQ-4279 (PDB: 9FCI)
USP7 + FT671 (PDB: 5NGE)
USP7 + GNE-6778 (PDB: 5UQX)
USP14 + IU1-248 (PDB: 6IIN)
USP28 + FT206 (PDB: 8P1Q)
USP30 + NK036 (PDB: 9F19)

**Fig. S3. Comparison of binding sites of selective USP DUB inhibitors.**

(**A**) USP9X catalytic domain (PDB: 5WCH (Paudel *et al*, 2019)) with the WEHI-092 binding site (6 aa motif: YVKGDL) highlighted in blue and catalytic triad in yellow. (**B**) Side by side comparison of co-crystal structures of USP DUBs and their selective inhibitors shown in red. The following co-crystal structures are visualised: USP1 in complex with KSQ-4279 (PDB: 9FCI (Cadzow *et al*, 2024; Rennie *et al*, 2024)), USP7 in complex with FT671 (PDB: 5NGE (Turnbull *et al*, 2017)), USP7 in complex with GNE-6778 (PDB: 5UQX (Kategaya *et al*, 2017)), USP14 in complex with IU1-248 (PDB: 6IIN (Wang *et al*, 2018)), USP28 in complex with FT206 (PDB: 8P1Q (Ruiz *et al*, 2021; Patzke *et al*, 2024)), USP30 in complex with NK036 (PDB: 9F19 (Kazi *et al*, 2025)).

### Figure S4

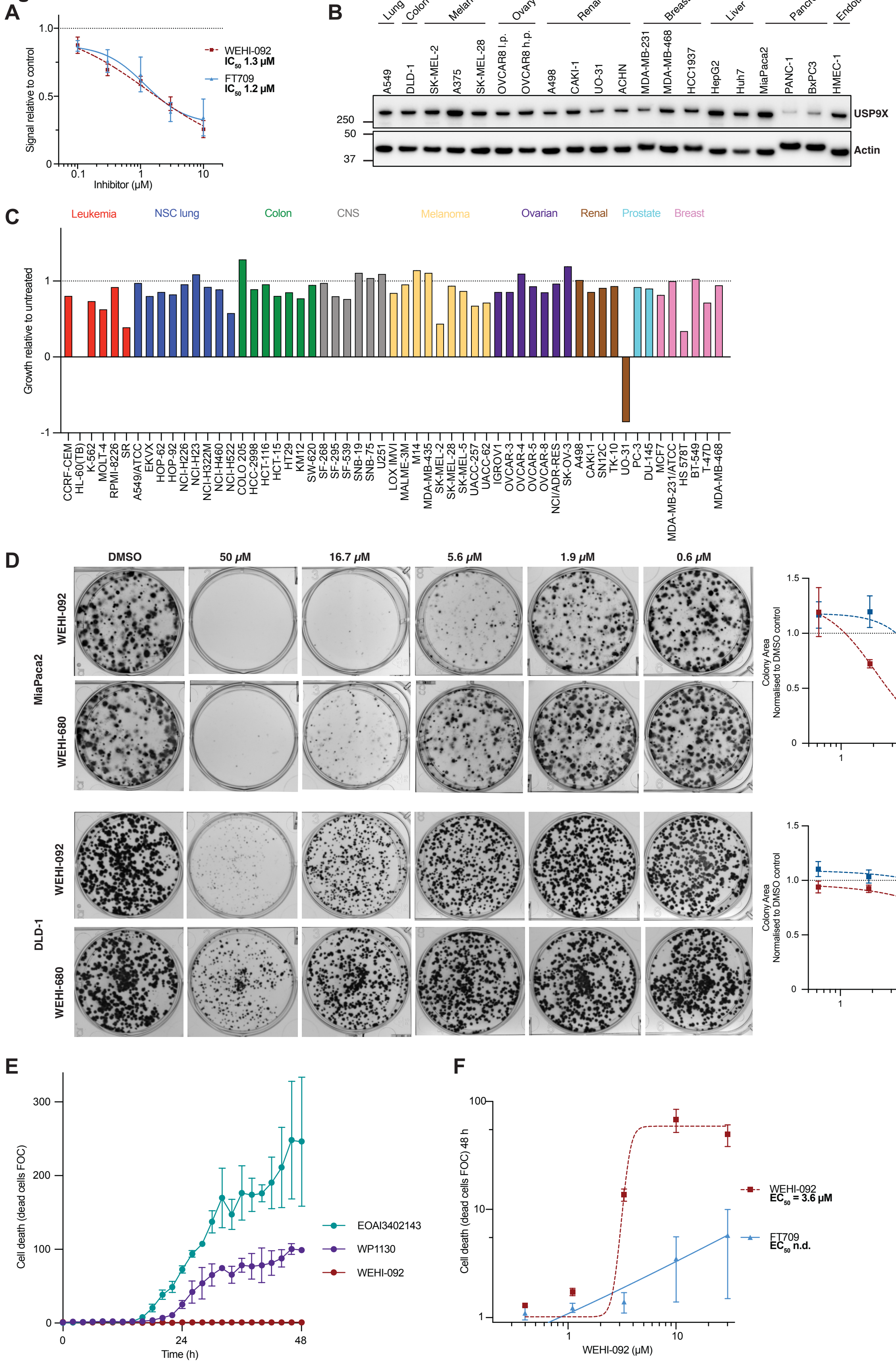
A
Signal relative to control
Inhibitor (μM)
WEHI-092
IC50 1.3 μM
FT709
IC50 1.2 μM
B
Lung
Colon
Melanoma
Ovary
Renal
Breast
Liver
Pancreas
Endothelial
A549
DLD-1
SK-MEL-2
A375
SK-MEL-28
OVCAR8 l.p.
OVCAR8 h.p.
A498
CAKI-1
UO-31
ACHN
MDA-MB-231
MDA-MB-468
HCC1937
HepG2
Huh7
MiaPaca2
PANC-1
BxPC3
HMEC-1
250
50
37
USP9X
Actin
C
Leukemia
NSC lung
Colon
CNS
Melanoma
Ovarian
Renal
Prostate
Breast
Growth relative to untreated
CCRF-CEM
HL-60(TB)
K-562
MOLT-4
RPMI-8226
SR
A549/ATCC
EKVX
HOP-62
HOP-92
NCI-H226
NCI-H23
NCI-H322M
NCI-H460
NCI-H522
COLO 205
HCC-2998
HCT-116
HCT-15
HT29
KM12
SW-620
SF-268
SF-295
SF-539
SNB-19
SNB-75
U251
LOX IMVI
MALME-3M
M14
MDA-MB-435
SK-MEL-2
SK-MEL-28
SK-MEL-5
UACC-257
UACC-62
IGROV1
OVCAR-3
OVCAR-4
OVCAR-5
OVCAR-8
NCI/ADR-RES
SK-OV-3
A498
CAKI-1
SN12C
TK-10
UO-31
PC-3
DU-145
MCF7
MDA-MB-231/ATCC
HS 578T
BT-549
T-47D
MDA-MB-468
D
DMSO
50 μM
16.7 μM
5.6 μM
1.9 μM
0.6 μM
MiaPaca2
DLD-1
WEHI-092
WEHI-680
MiaPaca2
Colony Area
Normalised to DMSO control
USP9Xi (μM)
WEHI-092
IC50 2.1 μM
WEHI-680
IC50 6.8 μM
DLD-1
E
Cell death (dead cells FOC)
Time (h)
EOAI3402143
WP1130
WEHI-092
F
Cell death (dead cells FOC) 48 h
WEHI-092 (μM)
WEHI-092
EC50 = 3.6 μM
FT709
EC50 n.d.

**Fig. S4. WEHI-092 phenotypes across cell lines.**

(**A**) $IC_{50}$ of USP9X inhibitors in MDA-MB-231 cells measured via degradation of the USP9X substrate CEP55 by Western Blot. The compounds were incubated with cells for 24 h. Data shown is the mean of three (WEHI-092) and two (FT709) independent biological repeats. Error bars, SEM. (**B**) USP9X expression levels by Western Blot across cell lines used in this study. l.p. - low passage; h.p. - high passage. Representative blot from two independent biological repeats is shown. (**C**) NCI-60 cancer cell panel of 57 cancer cell lines treated with 10 μM WEHI-680 for 3 days. Data shown is the cell growth at 3 days post treatment relative to untreated control and relative to day 0 for each individual cell line. Negative values indicate cell death at 3 days post treatment. Cancer cell lines are colour-coded depending on the cancer type. NCS - non-small cell. CNS - central nervous system. Data shown is the mean of two technical replicates. (**D**) WEHI-092 and WEHI-680 titration of clonogenic potential assays for MiaPaca2 and DLD-1 cells. *Left* Representative images from three biological repeats with three technical replicates each. *Right* Quantification of images plotted as dose-dependent colony area. Curve shown is the nonlinear curve fit generated using GraphPad Prism (v10.3) and was used for calculating the $IC_{50}$ values. Error bars, SEM. (**E**) Incucyte live cell imaging data for primary human dermal fibroblast cells. Data points shown are the mean of two independent biological repetitions with two technical replicates each. FOC - fold over control. EOAI3402143, WP1130 and WEHI-092 were used at 50 μM. Error bars, SEM. (**F**) Incucyte live cell imaging data showing the effects of WEHI-092 and FT709 on cell killing of UO-31 cells. Data points shown are the mean of three biological repetitions with two technical replicates each. Curve shown is the nonlinear curve fit generated using

GraphPad Prism (v10.3) and was used for calculating the $EC_{50}$ (n.d. – not determined). FOC - fold over control. Error bars, SEM.

Figure S5

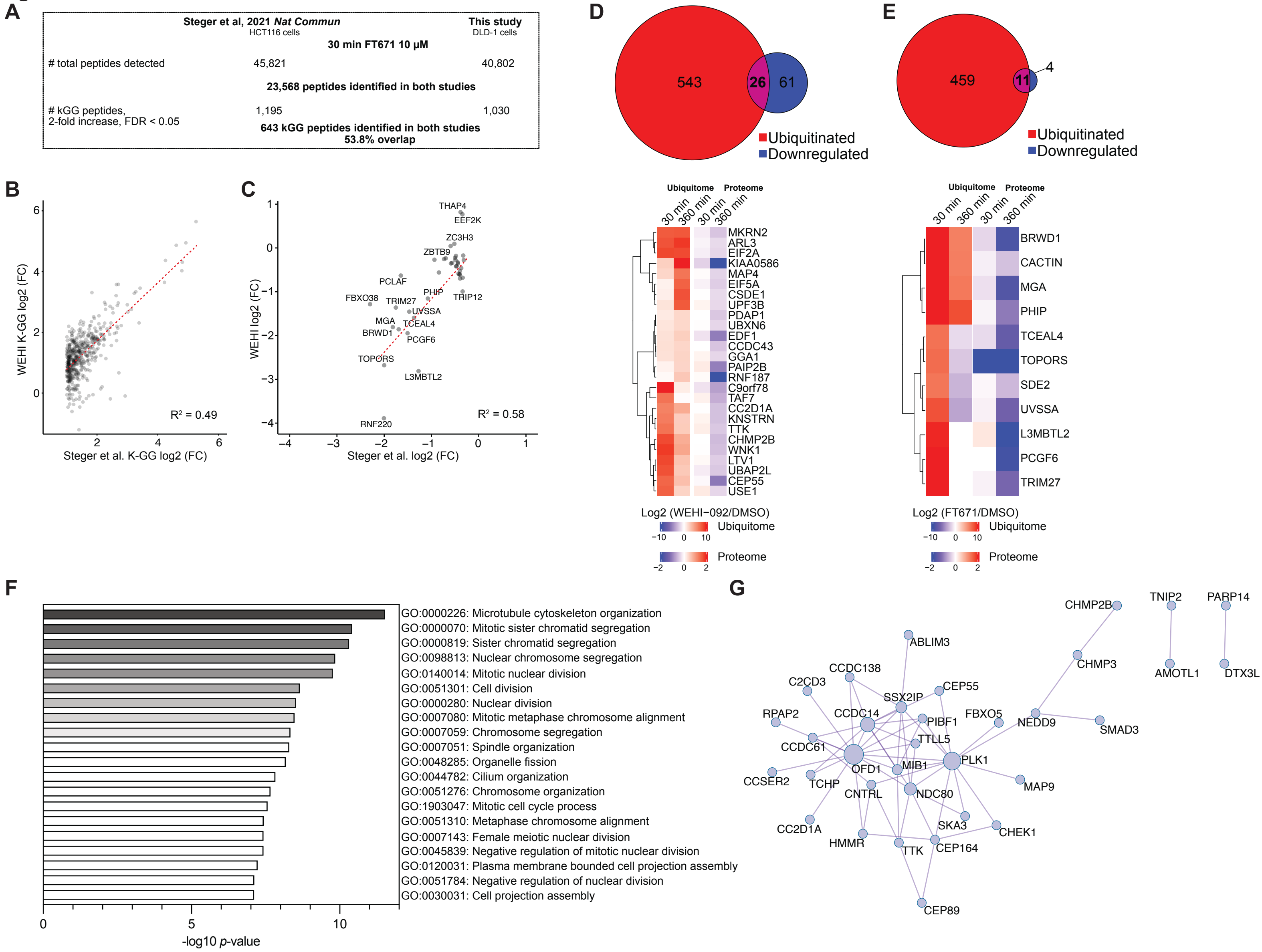
A
Steger et al, 2021 Nat Commun
HCT116 cells
This study
DLD-1 cells
30 min FT671 10 µM
### total peptides detected
45,821
40,802
23,568 peptides identified in both studies
### kGG peptides,
2-fold increase, FDR < 0.05
1,195
1,030
643 kGG peptides identified in both studies
53.8% overlap
B
WEHI K-GG log2 (FC)
Steger et al. K-GG log2 (FC)
R² = 0.49
C
WEHI log2 (FC)
Steger et al. log2 (FC)
R² = 0.58
THAP4
EEF2K
ZC3H3
ZBTB9
PCLAF
PHIP
TRIP12
FBXO38
TRIM27
UVSSA
MGA
TCEAL4
BRWD1
PCGF6
TOPORS
L3MBTL2
RNF220
D
543
26
61
Ubiquitinated
Downregulated
Ubiquitome
Proteome
30 min
360 min
30 min
360 min
MKRN2
ARL3
EIF2A
KIAA0586
MAP4
EIF5A
CSDE1
UPF3B
PDAP1
UBXN6
EDF1
CCDC43
GGA1
PAIP2B
RNF187
C9orf78
TAF7
CC2D1A
KNSTRN
TTK
CHMP2B
WNK1
LTV1
UBAP2L
CEP55
USE1
Log2 (WEHI−092/DMSO)
Ubiquitome
Proteome
E
459
11
4
Ubiquitinated
Downregulated
Ubiquitome
Proteome
30 min
360 min
30 min
360 min
BRWD1
CACTIN
MGA
PHIP
TCEAL4
TOPORS
SDE2
UVSSA
L3MBTL2
PCGF6
TRIM27
Log2 (FT671/DMSO)
Ubiquitome
Proteome
F
GO:0000226: Microtubule cytoskeleton organization
GO:0000070: Mitotic sister chromatid segregation
GO:0000819: Sister chromatid segregation
GO:0098813: Nuclear chromosome segregation
GO:0140014: Mitotic nuclear division
GO:0051301: Cell division
GO:0000280: Nuclear division
GO:0007080: Mitotic metaphase chromosome alignment
GO:0007059: Chromosome segregation
GO:0007051: Spindle organization
GO:0048285: Organelle fission
GO:0044782: Cilium organization
GO:0051276: Chromosome organization
GO:1903047: Mitotic cell cycle process
GO:0051310: Metaphase chromosome alignment
GO:0007143: Female meiotic nuclear division
GO:0045839: Negative regulation of mitotic nuclear division
GO:0120031: Plasma membrane bounded cell projection assembly
GO:0051784: Negative regulation of nuclear division
GO:0030031: Cell projection assembly
-log10 p-value
G
CHMP2B
TNIP2
PARP14
ABLIM3
CHMP3
AMOTL1
DTX3L
CCDC138
C2CD3
CEP55
SSX2IP
RPAP2
CCDC14
PIBF1
FBXO5
NEDD9
SMAD3
TTLL5
CCDC61
PLK1
OFD1
MIB1
CCSER2
TCHP
MAP9
CNTRL
NDC80
CC2D1A
SKA3
CHEK1
HMMR
TTK
CEP164
CEP89

**Fig. S5. USP7i and USP9Xi ubiquitinomics data.**

(**A**) Comparison of data output numbers of ubiquitinomics experiments benchmarking our study against the study on USP7 inhibition by Steger and colleagues (Steger *et al*, 2021) using the USP7 inhibitor FT671 on colorectal carcinoma cell lines. (**B**) Comparison between significantly increased K-GG peptides at 30 min between the study on USP7 inhibition by Steger and colleagues (Steger *et al*, 2021) (filtered for log2 > 1-fold change FT671 over untreated control, FDR < 0.05) and our study (FDR < 0.05). The data was pre-filtered allowing a missingness of zero. (**C**) Comparison between proteins downregulated (log2 < -0.3-fold change FT671 over untreated control, FDR < 0.05) in the study on USP7 inhibition by Steger and colleagues (Steger *et al*, 2021) and our study. BPCA+MinD was used for imputation. (**D**) DLD-1 time-resolved proteome and ubiquitome profiling upon WEHI-092 treatment (10 µM) for 30 min and 360 min. The data was filtered for increased ubiquitination (log2 > 2-fold change over untreated control, FDR < 0.05) at any timepoint and protein level decrease (log2 < -0.3 fold change over untreated control, FDR < 0.05) at any timepoint. Heatmap colours indicate fold changes in protein ubiquitination (left) and protein expression (right). Hierarchical clustering was performed on proteins (rows) with Euclidean distance as the similarity metric. The Venn diagram shows the number of ubiquitinated proteins, and the number of proteins depleted at any timepoint including overlapping proteins which are labelled in the heatmap. (**E**) DLD-1 time-resolved proteome and ubiquitome profiling upon USP7 inhibitor treatment with FT671 (10 µM) for 30 min and 360 min. The data was filtered for increased ubiquitination (log2 > 2-fold change over untreated control, FDR < 0.05) at any timepoint and protein level decrease (log2 < -0.3 fold change over untreated control, FDR < 0.05) at any timepoint. Heatmap colours indicate fold changes in protein ubiquitination (left) and protein expression (right). Hierarchical clustering was performed on proteins (rows) with Euclidean distance as the similarity metric. The Venn diagram shows the number of ubiquitinated proteins, and the number of proteins depleted at any timepoint including overlapping proteins labelled in the heatmap. (**F**) Pathway and process enrichment analysis of ubiquitinomic and proteomic analysis using metascape.org (Zhou *et al*, 2019). The top 20 enriched gene ontology terms were quantitatively ranked in a bar plot by their log10 *p*-value. (**G**) Protein-protein interaction enrichment analysis of 69 high confidence USP9X substrates in MDA-MB-231 cells (from **Fig. 4E**) using metascape.org (Zhou *et al*, 2019). A subset of 34 out of the 69 high confidence substrates was mapped in a protein-protein interaction network. The size of the circles corresponds to the number of mapped interactions.

### Figure S6

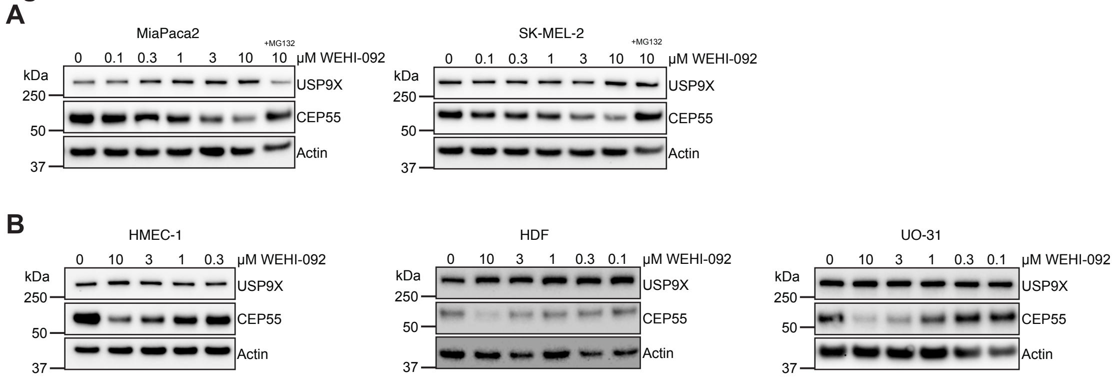

A
MiaPaca2
+MG132
0 0.1 0.3 1 3 10 10 µM WEHI-092
kDa
250
50
37
USP9X
CEP55
Actin
SK-MEL-2
+MG132
0 0.1 0.3 1 3 10 10 µM WEHI-092
kDa
250
50
37
USP9X
CEP55
Actin
B
HMEC-1
0 10 3 1 0.3 µM WEHI-092
kDa
250
50
37
USP9X
CEP55
Actin
HDF
0 10 3 1 0.3 0.1 µM WEHI-092
kDa
250
50
37
USP9X
CEP55
Actin
UO-31
0 10 3 1 0.3 0.1 µM WEHI-092
kDa
250
50
37
USP9X
CEP55
Actin

**Fig. S6. CEP55 protein levels are reduced in every cell line treated with WEHI-092.** (**A**) Western Blot validation of USP9X and CEP55 in MiaPaca2 and SK-MEL-2 cells upon WEHI-092 treatment (24 h). Proteasomal inhibition using MG132 was performed at 10 μM for 24 h in presence of 5 μM QVD apoptosis blockage. The blots shown are a representative of at least two biological repeats. (**B**) Western Blot validation of USP9X and CEP55 in HMEC-1, Human dermal fibroblasts (HDF) and UO-31 cells (24 h). The blots shown are a representative of at least two biological repeats.

### Figure S7

## A

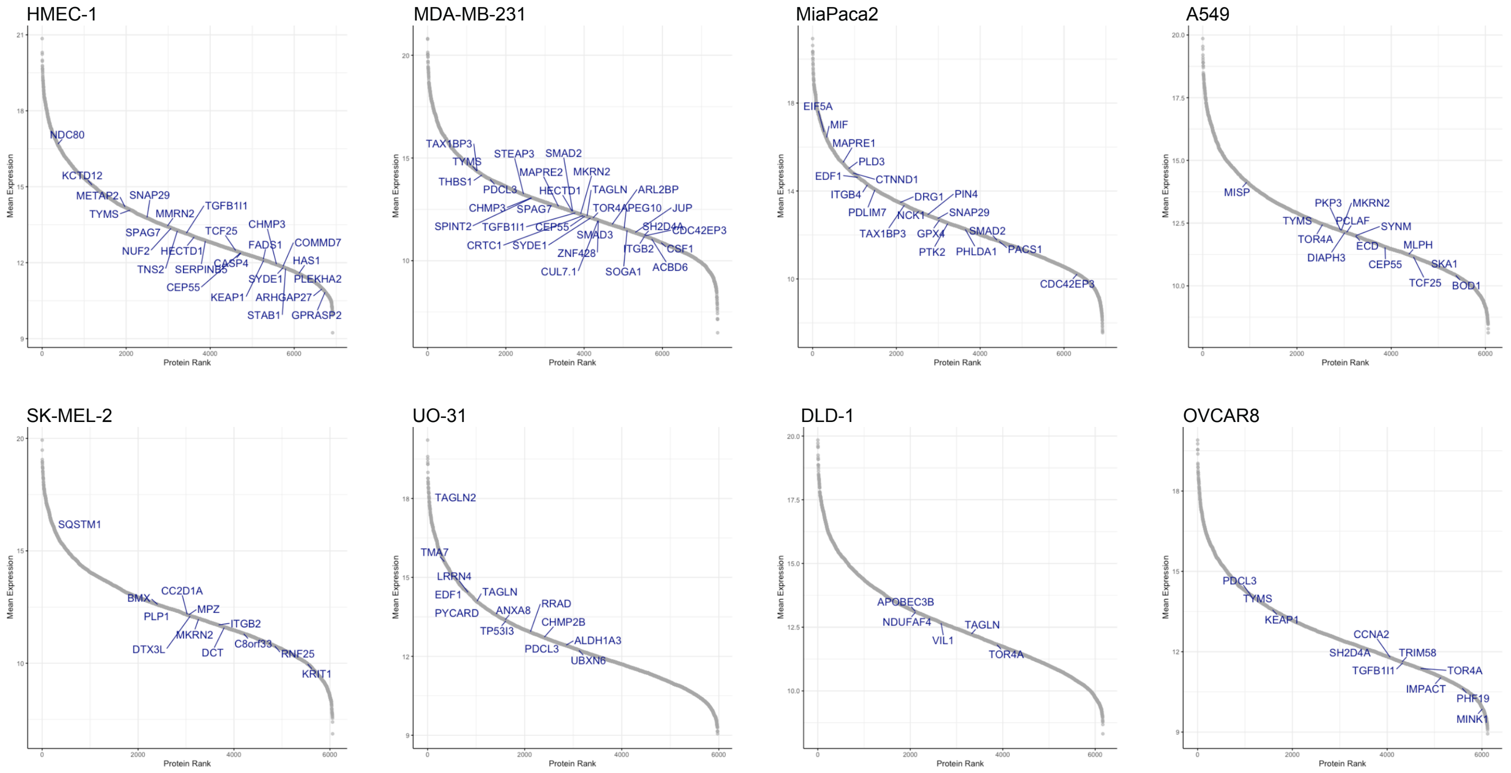

HMEC-1
NDC80
KCTD12
METAP2
SNAP29
TYMS
MMRN2
TGFB1I1
SPAG7
NUF2
HECTD1
TCF25
CHMP3
TNS2
SERPINB5
CASP4
FADS1
COMMD7
HAS1
CEP55
SYDE1
PLEKHA2
KEAP1
ARHGAP27
STAB1
GPRASP2
MDA-MB-231
TAX1BP3
STEAP3
SMAD2
TYMS
MAPRE2
MKRN2
THBS1
PDCL3
HECTD1
TAGLN
ARL2BP
CHMP3
SPAG7
TOR4A
PEG10
JUP
SPINT2
TGFB1I1
CEP55
SH2D4A
CDC42EP3
SMAD3
CRTC1
SYDE1
ITGB2
CSF1
ZNF428
CUL7.1
SOGA1
ACBD6
MiaPaca2
EIF5A
MIF
MAPRE1
PLD3
EDF1
CTNND1
ITGB4
DRG1
PIN4
PDLIM7
NCK1
SNAP29
TAX1BP3
GPX4
SMAD2
PTK2
PHLDA1
PACS1
CDC42EP3
A549
MISP
PKP3
MKRN2
TYMS
PCLAF
SYNM
TOR4A
ECD
MLPH
DIAPH3
CEP55
SKA1
TCF25
BOD1
SK-MEL-2
SQSTM1
BMX
CC2D1A
PLP1
MPZ
ITGB2
MKRN2
DTX3L
DCT
C8orf33
RNF25
KRIT1
UO-31
TAGLN2
TMA7
LRRN4
EDF1
TAGLN
RRAD
PYCARD
ANXA8
CHMP2B
TP53I3
ALDH1A3
PDCL3
UBXN6
DLD-1
APOBEC3B
NDUFAF4
TAGLN
VIL1
TOR4A
OVCAR8
PDCL3
TYMS
KEAP1
CCNA2
SH2D4A
TRIM58
TGFB1I1
TOR4A
IMPACT
PHF19
MINK1
Mean Expression
Protein Rank

**Fig. S7. Proteins depleted upon WEHI-092 treatment vary in expression levels in the eight different cell lines used.**

(**A**) Proteins ranked by expression as a mean over 5 technical replicates (untreated control samples used). Proteins found to be depleted upon WEHI-092 treatment (**Fig. 6A**) are labelled (filtered for log2 <-0.585 fold change over untreated control condition, p.adj < 0.05) in each cell line shown.

### Figure S8

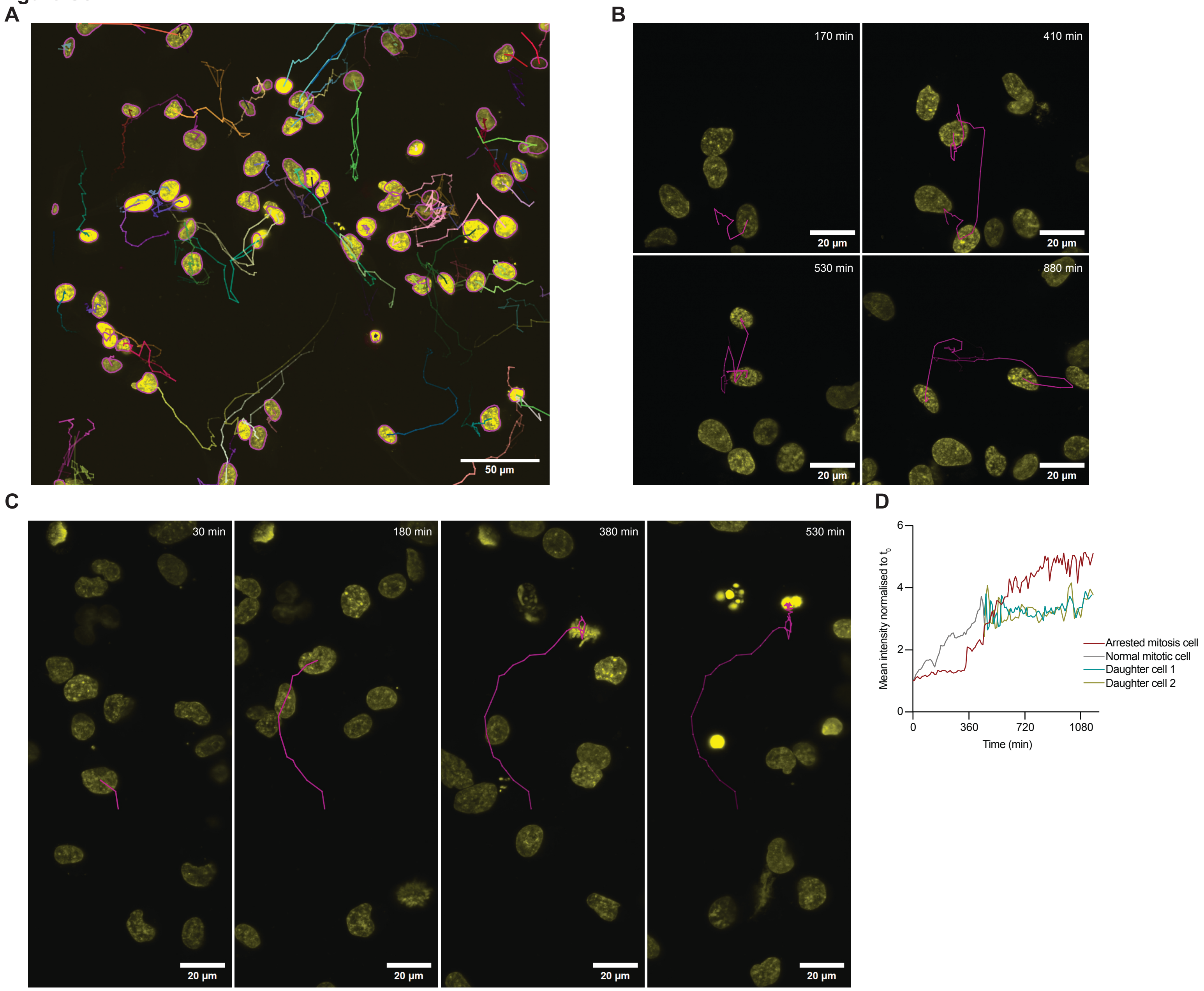

A
50 µm
B
170 min
410 min
530 min
880 min
20 µm
C
30 min
180 min
380 min
530 min
20 µm
D
Mean intensity normalised to $t_0$
Time (min)
0
360
720
1080
2
4
6
Arrested mitosis cell
Normal mitotic cell
Daughter cell 1
Daughter cell 2

**Fig. S8. Lattice light sheet imaging analysis.**

(**A**) Demonstration of a full field of MDA-MB-231 cells with associated segmentation, via StarDist, and associated individual cell tracks, using an advanced Kalman tracker capable of merging and splitting tracks. Scale bar: 50 µm. (**B**) Example of a mitotic MDA-MB-231 cell (yellow) with its associated track (pink). Following mitosis, the track continues to follow the two daughter cells. Scale bar: 20 µm. (**C**) Example of mitotic arrest of MDA-MB-231 cell (yellow) with associated track (pink) over time left to right. Scale bar: 20 µm. (**D**) Plot of mean pixel intensity over time for normal mitotic cell from **B**, shown in a grey line and arrested mitosis cell from **C**, shown in a red line. Grey line depicts mean intensity for a single cell as it enters mitosis with the two green lines denoting the mean pixel intensity for each daughter cell. The data was normalized to $t_0$.

**Movie S1**

Movie S1 shows an example of a normal cell division in MDA-MB-231 cells over time. DNA is shown in yellow (right), and tubulin is shown in magenta (middle). Merged channels are shown on the left. Timestamp is in h. Scale bar: 20 µm.

**Movie S2**

Movie S2 shows an example of mitotic arrest in MDA-MB-231 cells over time. DNA is shown in yellow (middle), and tubulin is shown in magenta (right). Merged channels are shown on the left. Timestamp is in h. Scale bar: 20 µm.

**Movie S3**

Movie S3 shows an example of the full-field timelapse for MiaPaca2, MDA-MB-231 and HMEC-1 cells with untreated DMSO control cells on the left and WEHI-092 (15 µM) treated cells on the right. DNA is shown in yellow, tubulin is shown in magenta, and AnnexinV is shown in cyan. Timestamp is in h and 24 h window is shown. Scale bar: 50 µm.

**Movie S4**

Movie S4 shows examples of microtubule disassembly upon WEHI-092 (15 µM) treatment in MDA-MB-231 cells. Tubulin is shown in a Fire LUT generated with ImageJ/Fiji.

**Table S1. List of cell lines including culture conditions**

| Cell line | Growth media | Source |
|---|---|---|
| MDA-MB-231 | DMEM + 10% FBS | WEHI |
| MDA-MB-468 | RMPI + 10% FBS + 1X GlutaMAX + 5 µg/mL Insulin | WEHI |
| MCF-7 | DMEM + 10% FBS + 1X Glutamine | WEHI |
| A549 | DMEM + 10% FBS | WEHI |
| ACHN | RPMI + 10% FBS | WEHI |
| CAKI-1 | RPMI + 10% FBS | WEHI |
| A498 | RPMI + 10% FBS | WEHI |
| UO-31 | RPMI + 10% FBS | WEHI |
| MiaPaca2 | DMEM + 10% FBS + 2% horse serum | WEHI |
| PANC-1 | DMEM + 10% FBS | WEHI |
| BxPC3 | RPMI + 10% FBS | WEHI |
| DLD-1 | DMEM + 10% FBS + 1X GlutaMAX | WEHI |
| HMEC-1 | Endothelial Cell Growth Medium (Sigma, 211-500) | WEHI |
| HUVEC | Endothelial Cell Growth Medium (Sigma, 211-500) | StemCell Technologies (US), Female |
| Human dermal fibroblasts (HDF) | DMEM + 20% FBS | Lonza (Switzerland) (19TL178960) P1 58Y, Female |
| A375 | DMEM + 10% FBS | WEHI |
| SK-MEL-2 | RPMI + 10% FBS | WEHI |
| SK-MEL-28 | RPMI + 10% FBS | WEHI |
| Huh7 | DMEM + 10% FBS | WEHI |
| HepG2 | DMEM + 10% FBS | WEHI |
| OVCAR8 | DMEM + 10% FBS + 5 µg/mL Insulin + 0.05 µg/mL EGF + 1 µg/mL Hydrocortisone | WEHI |
| THP-1 | RPMI + 10% FBS | WEHI |

**Supplementary Material: Medicinal Chemistry**

All reagents were used as received from commercial suppliers unless stated otherwise. NMR spectra were recorded at ambient temperature on a Bruker Avance 300 MHz instrument in the specified deuterated solvent. Observed proton chemical shifts were reported as units of parts per million (ppm) relative to the respective residual solvent peak DMSO-d6 ($\delta$ 2.50). Multiplicities were reported: s (singlet), d (doublet), t (triplet), q (quartet), dd (doublet of doublets), and m (multiplet). Coupling constants were reported as a *J* value in Hertz (Hz). Abbreviations: DCM (dichloromethane), DMF (N,N-dimethylformamide), DIPEA (N,N-diisopropylethylamine), EDCI (*N*-(3-dimethylaminopropyl)-*N′*-ethylcarbodiimide), 1-hydroxybenzotriazole (HOBt) and MeOH (methanol).

Commercial compounds including FT671, FT709, WP1130, EOAI3402143 and PR619 were sourced from MedChemExpress. The negative control compound WEHI-871 (3-[4-(2,3-dihydro-1,4-benzodioxine-6-sulfonylpiperazin-1-yl]-6-phenylpyridazine) was sourced from Enamine.

**Synthetic procedures**

Synthesis of (4-((2,3-dihydrobenzo[*b*][1,4]dioxin-6-yl)sulfonyl)piperazin-1-yl)(2,3-dimethyl-1*H*-indol-5-yl)methanone **(WEHI-092)**, (4-((2,3-dihydrobenzo[*b*][1,4]dioxin-6-yl)sulfonyl)piperazin-1-yl)(1,2,3-trimethyl-1*H*-indol-5-yl)methanone **(WEHI-333)**, (4-((2,3-dihydrobenzo[*b*][1,4]dioxin-6-yl)sulfonyl)piperazin-1-yl)(1*H*-indol-5-yl)methanone **(WEHI-680)** and 6-(4-((2,3-dihydrobenzo[*b*][1,4]dioxin-6-yl)sulfonyl)piperazine-1-carbonyl)quinoxaline-2,3(1*H*,4*H*)-dione **(WEHI-553) (see Fig. S1A)**

(4-((2,3-Dihydrobenzo[*b*][1,4]dioxin-6-yl)sulfonyl)piperazin-1-yl)(2,3-dimethyl-1*H*-indol-5-yl)methanone **(WEHI-092)**

To a stirred solution of 2,3-dimethyl-1*H*-indole-5-carboxylic acid (206 mg, 1.09 mmol), HOBt hydrate (184 mg, 1.20 mmol) and EDCI hydrochloride (251 mg, 1.31 mmol) in DMF (5 mL) was added DIPEA (665 µL, 3.82 mmol) and 1-((2,3-dihydrobenzo[*b*][1,4]dioxin-6-yl)sulfonyl)piperazine hydrochloride (350 mg, 1.09 mmol). The solution was stirred at ambient temperature for 16h. The solution was poured onto ice/water (200 mL), whereby a precipitate formed. The solid was filtered and washed with water to give crude product (487 mg). This was purified by column chromatography (10% MeOH in DCM), fractions combined, and solvent evaporated under reduced pressure. Water was added and the whole stirred, filtered and washed with water to give 4-((2,3-dihydrobenzo[*b*][1,4]dioxin-6-yl)sulfonyl)piperazin-1-yl)(2,3-dimethyl-1*H*-indol-5-yl)methanone as a beige solid (431 mg, 87%). $^1$H NMR (300 MHz, DMSO) δ

10.89 (s, 1H), 7.39 (s, 1H), 7.23 – 7.15 (m, 3H), 7.10 (d, *J* = 8.3 Hz, 1H), 6.97 (dd, *J* = 8.3, 1.6 Hz, 1H), 4.34 (q, *J* = 5.2 Hz, 4H), 3.65 – 3.55 (m, 4H), 2.96 – 2.86 (m, 4H), 2.30 (s, 3H), 2.13 (s, 3H). $^{13}$C NMR (75 MHz, DMSO) $\delta$ 171.1, 147.6, 143.5, 135.7, 132.8, 128.3, 127.1, 124.7, 121.1, 119.4, 117.8, 117.2, 116.5, 109.7, 105.8, 64.4, 64.0, 45.9, 11.2, 8.2.

(4-((2,3-dihydrobenzo[*b*][1,4]dioxin-6-yl)sulfonyl)piperazin-1-yl)(1,2,3-trimethyl-1*H*-indol-5-yl)methanone (**WEHI-033)**

To a stirred solution of 1,2,3-trimethylindole-5-carboxylic acid (33 mg, 0.162 mmol), HOBt hydrate (26 mg, 0.170 mmol) and EDCI hydrochloride (36 mg, 0.188 mmol) in DMF (1 mL) was added DIPEA (95 µL, 0.546 mmol) and 1-((2,3-dihydrobenzo[*b*][1,4]dioxin-6-yl)sulfonyl)piperazine hydrochloride (50 mg, 0.156 mmol). The solution was stirred at ambient temperature for 16h. The solution was poured onto ice/water (50 mL) and stirred at ambient temperature for 2h, whereby a precipitate formed. The solid was filtered, washed with water to give (4-((2,3-dihydrobenzo[*b*][1,4]dioxin-6-yl)sulfonyl)piperazin-1-yl)(1,2,3-trimethyl-1*H*-indol-5-yl)methanone as a beige solid (58 mg, 79%). $^{1}$H NMR (300 MHz, DMSO) $\delta$ 7.43 (d, *J* = 1.5 Hz, 1H), 7.34 (d, *J* = 8.4 Hz, 1H), 7.23 – 7.15 (m, 2H), 7.10 (d, *J* = 8.3 Hz, 1H), 7.05 (dd, *J* = 8.4, 1.6 Hz, 1H), 4.34 (q, *J* = 4.7 Hz, 4H), 3.64 (s, 3H), 3.59 (br s, 4H), 2.92 (br s, 4H), 2.33 (s, 3H), 2.17 (s, 3H). $^{13}$C NMR (75 MHz, DMSO) $\delta$ 171.0, 147.6, 143.5, 136.7, 134.5, 127.3, 127.2, 124.9, 121.1, 119.5, 117.8, 117.4, 116.5, 108.4, 105.9, 64.4, 64.0, 45.9, 29.5, 9.8, 8.5.

(4-((2,3-dihydrobenzo[*b*][1,4]dioxin-6-yl)sulfonyl)piperazin-1-yl)(1*H*-indol-5-yl)methanone **(WEHI-680)**

To a stirred solution of 1*H*-indole-5-carboxylic acid (26 mg, 0.161 mmol), HOBt hydrate (26 mg, 0.170 mmol) and EDCI hydrochloride (36 mg, 0.188 mmol) in DMF (1 mL) was added DIPEA (95 µL, 0.546 mmol) and 1-((2,3-dihydrobenzo[*b*][1,4]dioxin-6-yl)sulfonyl)piperazine hydrochloride (50 mg, 0.156 mmol). The solution was stirred at ambient temperature for 16h. The solution was poured onto ice/water (50 mL) and stirred at ambient temperature for 2h, whereby a precipitate formed. The solid was filtered, washed with water to give (4-((2,3-dihydrobenzo[*b*][1,4]dioxin-6-yl)sulfonyl)piperazin-1-yl)(1*H*-indol-5-yl)methanone as a beige solid (64.0 mg, 96%). $^{1}$H NMR (300 MHz, DMSO) $\delta$ 11.29 (s, 1H), 7.58 (d, $J$ = 1.5 Hz, 1H), 7.44 – 7.36 (m, 2H), 7.24 – 7.16 (m, 2H), 7.14 – 7.05 (m, 2H), 6.47 (t, $J$ = 2.4 Hz, 1H), 4.34 (q, $J$ = 4.9 Hz, 4H), 3.60 (s, 4H), 2.92 (t, $J$ = 4.9 Hz, 4H). $^{13}$C NMR (75 MHz, DMSO) $\delta$ 170.8, 147.6, 143.5, 136.4, 127.2, 126.9, 126.6, 125.6, 121.1, 120.5, 119.8, 117.8, 116.5, 111.1, 101.8, 64.4, 64.0, 45.9.

6-(4-((2,3-dihydrobenzo[*b*][1,4]dioxin-6-yl)sulfonyl)piperazine-1-carbonyl)quinoxaline-2,3(1*H*,4*H*)-dione (**WEHI-553**)

To a stirred solution of 2,3-dioxo-1,4-dihydroquinoxaline-6-carboxylic acid hydrate (35 mg, 0.156 mmol), HOBt hydrate (26 mg, 0.170 mmol) and EDCI hydrochloride (35 mg, 0.183 mmol) in DMF (1 mL) was added DIPEA (95 µL, 0.546 mmol) and 1-((2,3-

dihydrobenzo[*b*][1,4]dioxin-6-yl)sulfonyl)piperazine hydrochloride (50 mg, 0.156 mmol). The solution was stirred at ambient temperature for 16h. The solution was poured onto ice/water (50 mL) and stirred at ambient temperature for 2h, whereby a white precipitate formed. The solid was filtered, washed with water to give 6-(4-((2,3-dihydrobenzo[*b*][1,4]dioxin-6-yl)sulfonyl)piperazine-1-carbonyl)quinoxaline-2,3(1*H*,4*H*)-dione as a fawn solid (44.2 mg, 60%). $^{1}$H NMR (300 MHz, DMSO) $\delta$ 12.00 (s, 2H), 7.23 – 7.07 (m, 6H), 4.34 (d, J = 4.3 Hz, 4H), 3.55 (s, 4H), 2.93 (s, 4H). $^{13}$C NMR (75 MHz, DMSO) $\delta$ 168.6, 155.2, 147.7, 143.5, 129.3, 127.2, 125.7, 122.1, 121.1, 117.9, 116.5, 115.0, 114.6, 64.4, 64.1, 45.8.

#### NMR Data

$^1H$ NMR of WEHI-092

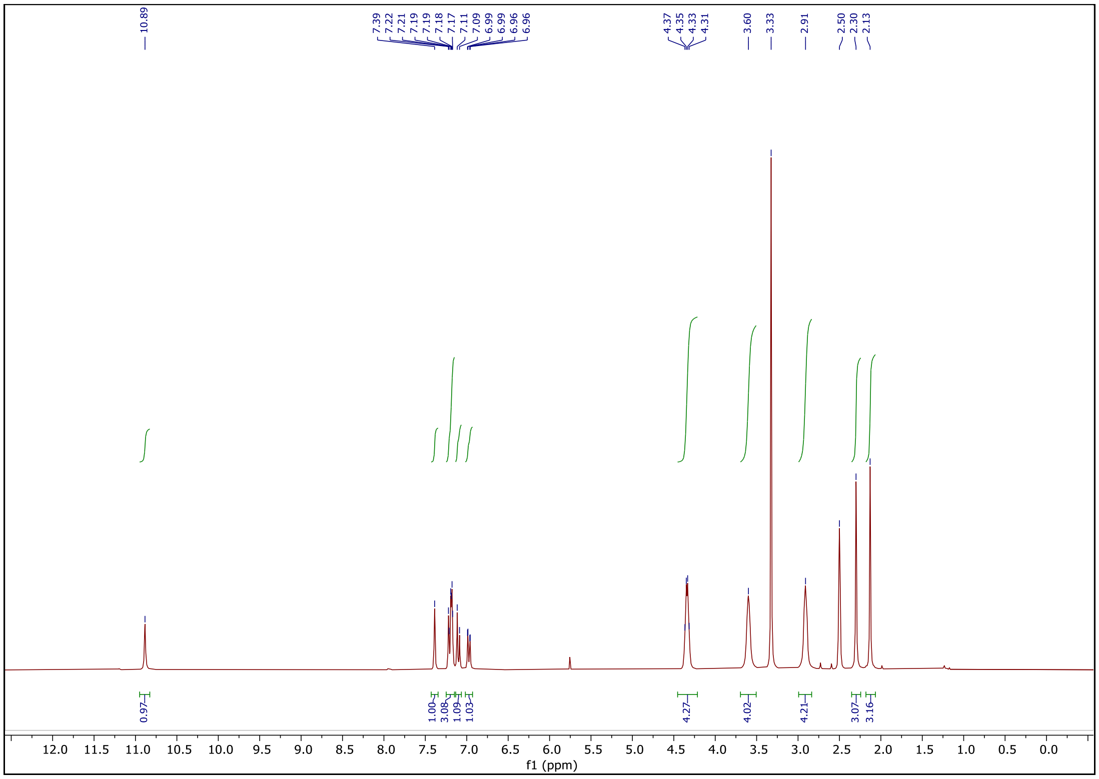


$^{13}C$ NMR of WEHI-092

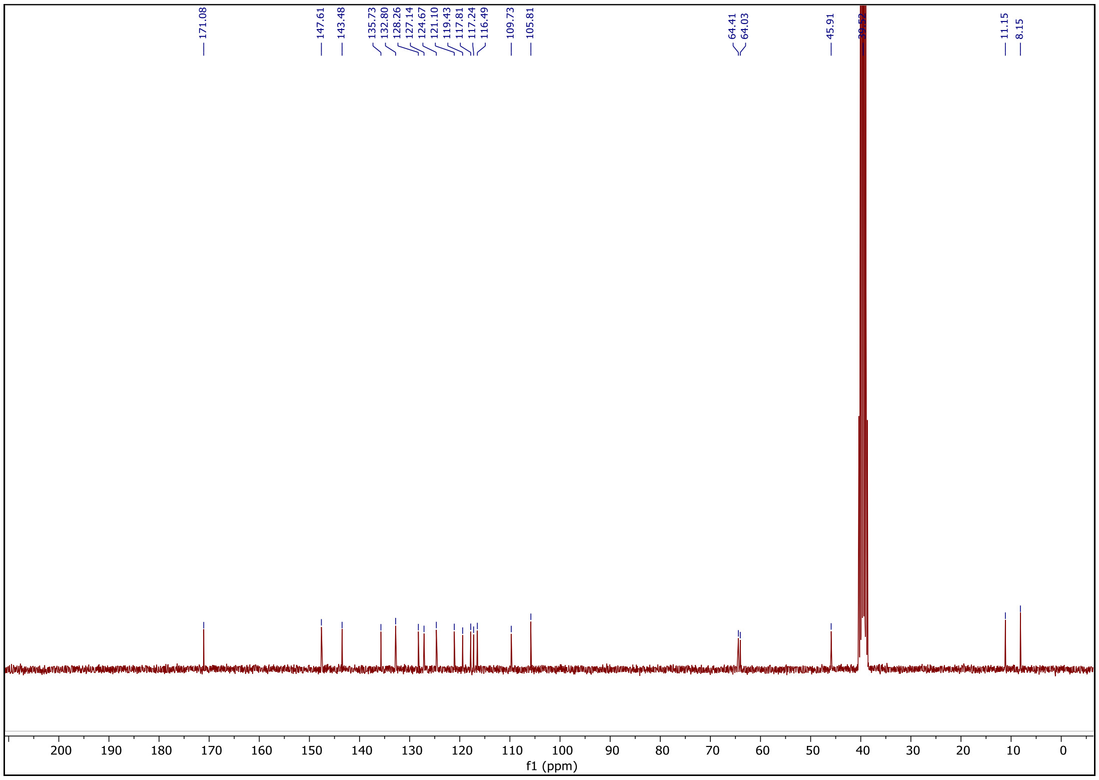

#### $^{1}H$ NMR of WEHI-333

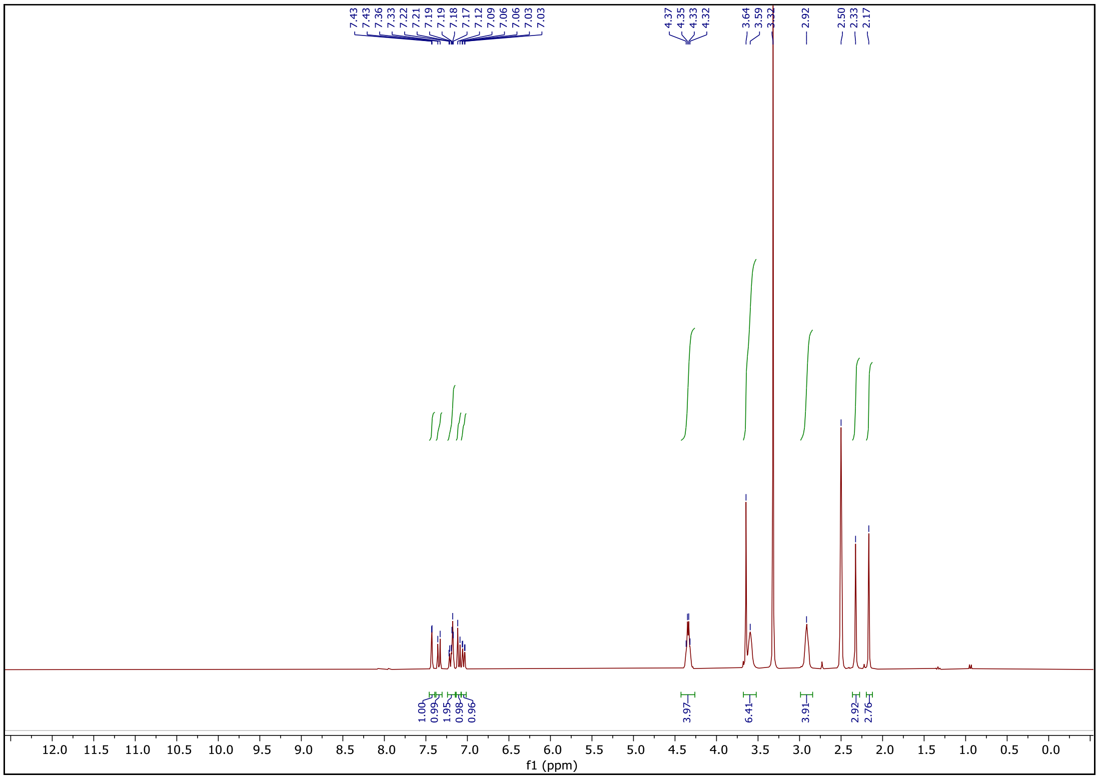


#### $^{13}C$ NMR of WEHI-333

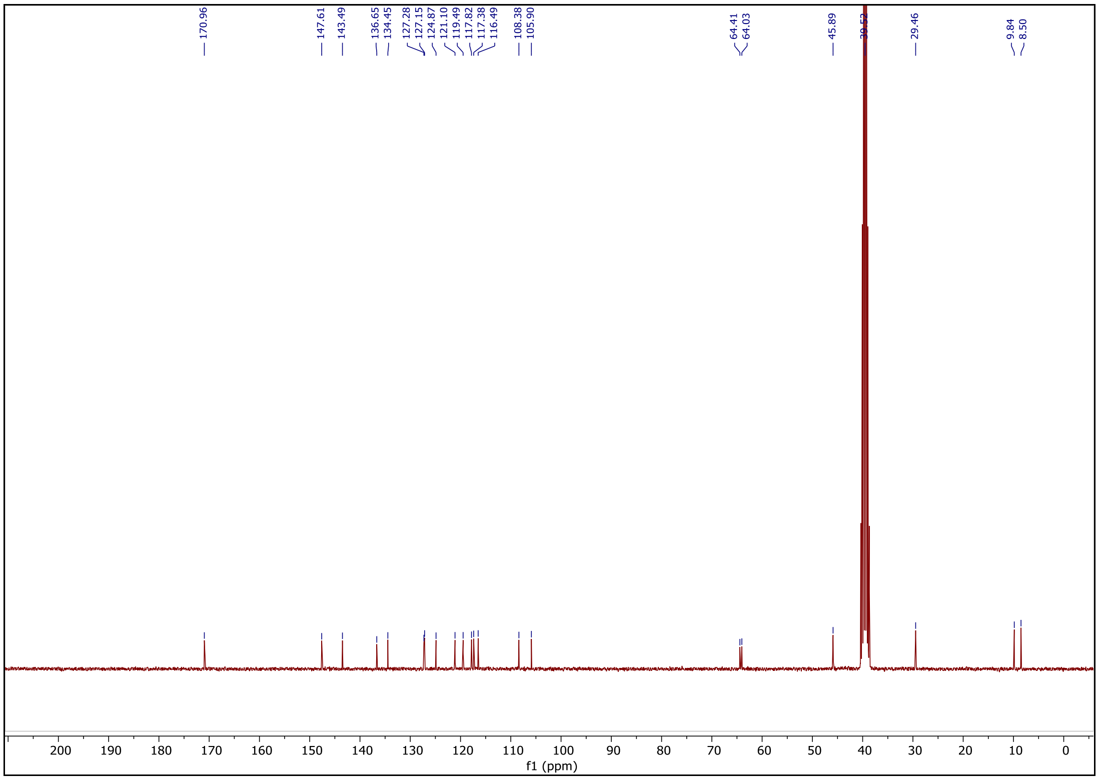

#### $^{1}$H NMR of WEHI-680

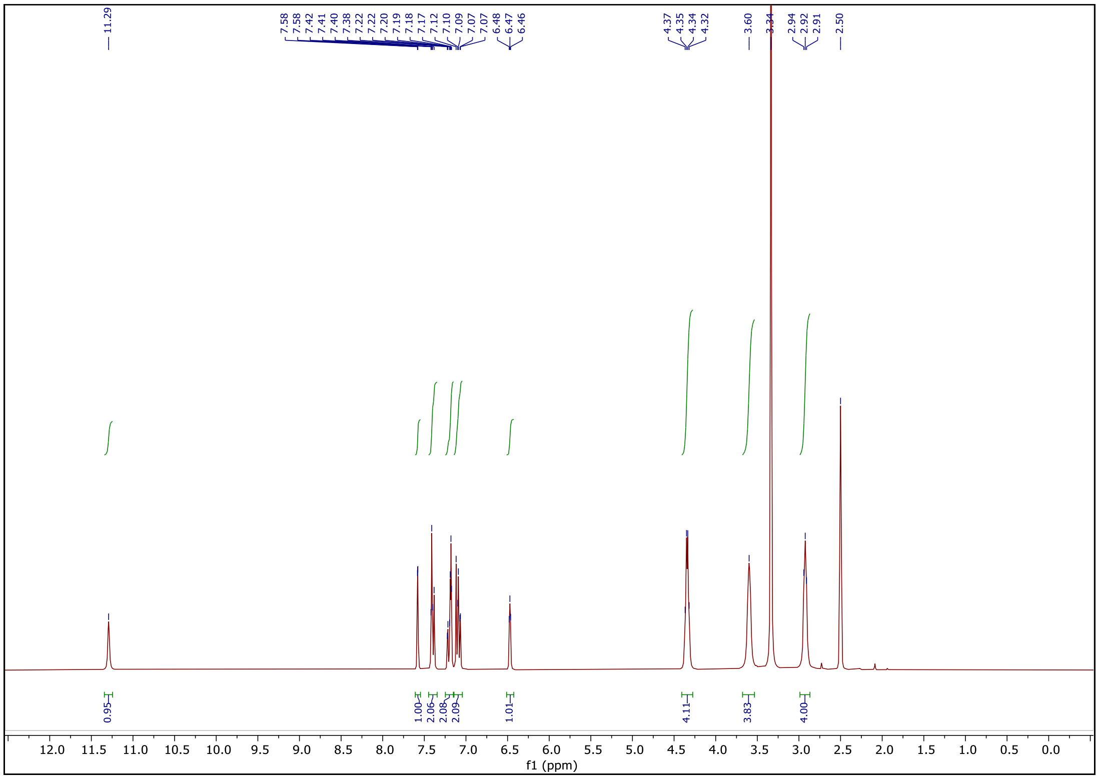


#### $^{13}$C NMR of WEHI-680

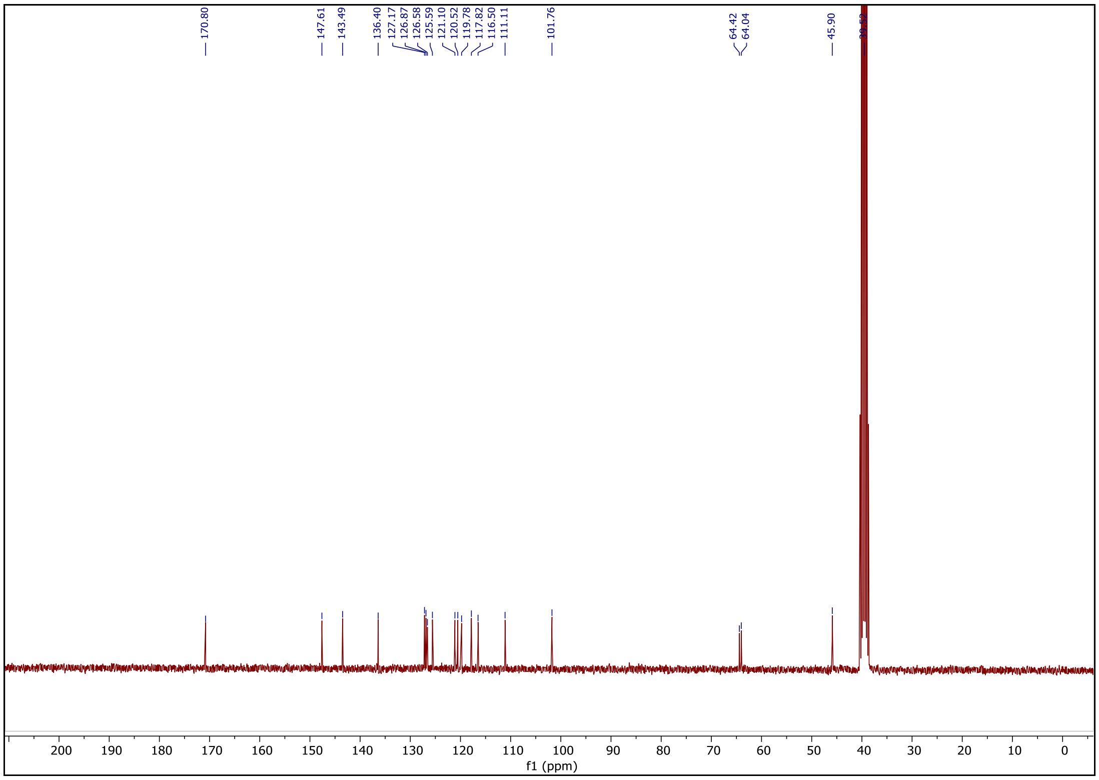

#### $^{1}$H NMR of WEHI-553

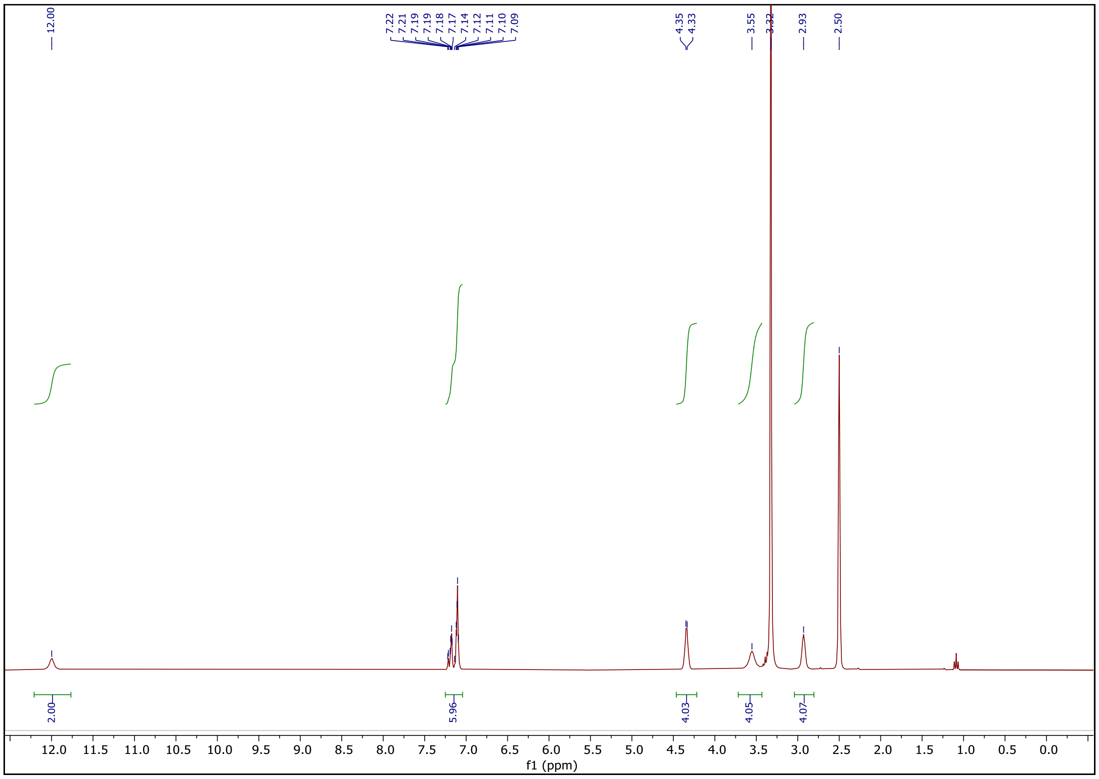


#### $^{13}$C NMR of WEHI-553

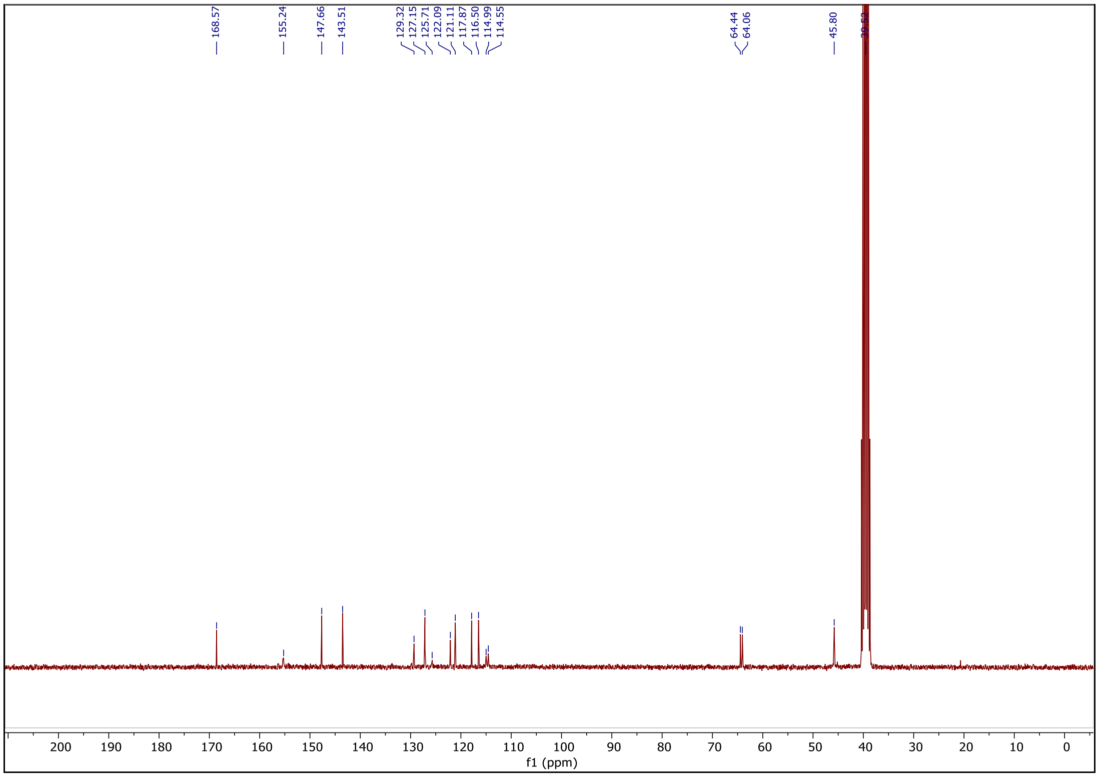
